# Degradation of the repetitive genomic landscape in a close relative of *C. elegans*

**DOI:** 10.1101/797035

**Authors:** Gavin C. Woodruff, Anastasia A. Teterina

**Affiliations:** Institute of Ecology and Evolution, University of Oregon, Eugene, OR, USA; Center of Parasitology, Severtsov Institute of Ecology and Evolution RAS, Moscow, Russia

## Abstract

*The abundance, diversity, and genomic distribution of repetitive elements is highly variable among species. These patterns are thought to be driven in part by reproductive mode and the interaction of selection and recombination, and recombination rates typically vary by chromosomal position. In the nematode* C. elegans, *repetitive elements are enriched at chromosome arms and depleted on centers, and this mirrors the chromosomal distributions of other genomic features such as recombination rate. How conserved is this genomic landscape of repeats, and what evolutionary forces maintain it? To address this, we compared the genomic organization of repetitive elements across five* Caenorhabditis *species with chromosome-level assemblies. As previously reported, repeat content is enriched on chromosome arms in most* Caenorhabditis *species, and no obvious patterns of repeat content associated with reproductive mode were observed. However, the fig-associated* Caenorhabditis inopinata *has experienced rampant repetitive element expansion and reveals no association of global repeat content with chromosome position. Patterns of transposable element superfamily-specific distributions reveal this global pattern is driven largely by a few transposable element superfamilies that in* C. inopinata *have expanded in number and have weak associations with chromosome position. Additionally, 15% of predicted protein-coding genes in* C. inopinata *align to transposon-related proteins. When these are excluded*, C. inopinata *has no enrichment of genes in chromosome centers, in contrast to its close relatives who all have such clusters. Forward evolutionary simulations reveal that chromosomal heterogeneity in recombination rate is insufficient for generating structured genomic repetitive landscapes. Instead, heterogeneity in the fitness effects of transposable element insertion is needed to promote heterogeneity in repetitive landscapes. Thus, patterns of gene density along chromosomes are likely drivers of global repetitive landscapes in this group, although other historical or genomic factors are needed to explain the idiosyncrasy of genomic organization of various transposable element taxa within* C. inopinata. *Taken together, these results highlight the power of comparative genomics and evolutionary simulations in testing hypotheses regarding the causes of genome organization.*

## Introduction

**R**epetitive elements are a conspicuous feature of eukaryotic genomes. Over half of the human genome is comprised of such elements (de Koning, et al. 2011), and the maize genome has a repeat content of over 80% (Baucom, et al. 2009; Schnable, et al. 2009). But at the same time, the range in repeat content among eukaryotic genomes is great, with some species having a scant number of repetitive elements (0.8% in one species of bdelloid rotifer (Nowell, et al. 2018)). And not only does the global repeat content among genomes vary—heterogeneity in repeat content both within and between chromosomes occurs (Rizzon, et al. 2002; Stitzer, et al. 2019). Furthermore, repeat density in genomes has been observed to covary with patterns of genomic diversity (Clark, et al. 2007), recombination rate (Rizzon, et al. 2002), gene density (Medstrand, et al. 2002), chromatin state (Peng and Karpen 2008), centromeric regions (Plohl, et al. 2014) and physical spatial position (Guelen, et al. 2008). Repetitive elements are then a major feature of genomic organization, and the origin and maintenance of their genomic landscape demands explanation.

Transposable elements are generally considered deleterious by replicating at the expense of its host and abrogating functional sites through insertion. This is largely consistent with experiments revealing that fitness declines with increased transposable element activity (Pasyukova, et al. 2004; Bégin and Schoen 2006). Thus it has been proposed that variation in repeat content among species is driven by variation in population size; weaker selection in species with smaller population sizes should lead to increased repeat content (Lynch 2007). This is also thought to explain within-genome heterogeneity in repeat content—low recombining regions have higher repeat content than regions with high recombination rates in multiple systems (including fruit flies (Rizzon, et al. 2002); yeast (Pan, et al. 2011); mice (Jensen-Seaman, et al. 2004); humans (Jensen-Seaman, et al. 2004); *Arabidopsis thaliana* (Wright, et al. 2003; Kent, et al. 2017); and maize (Liu, et al. 2009), among others). This is presumably because of weakened selection in low-recombining regions (Hill and Robertson 1966; Langley, et al. 1988). However, in mice and humans, the association of transposable element content with recombination rate varies depending on repeat type (Myers, et al. 2005; Shifman, et al. 2006). Furthermore, transposable element activity was not correlated with recombination rate after accounting for chromatin states in *Drosophila melanogaster* (Adrion, et al. 2017). Others have noted that transposable element abundance should either increase or decrease depending on the model of natural selection and the mode of reproduction (Wright and Schoen 1999; Bestor 2000; Morgan 2001; Glémin, et al. 2019). At the same time, it has also been proposed that repetitive elements themselves can be adaptive (Shapiro and Von Sternberg 2005), and individual cases of adaptation via transposable element insertion abound (Oliver and Greene 2009; Casacuberta and González 2013). And in nematodes, repeat content is atypically positively correlated with recombination (Duret, et al. 2000), and little difference is observed between selfing and outcrossing close relatives (Fierst, et al. 2015). What drives the landscape of repetitive elements both between species and within genomes?

One approach towards tackling this question is investigating a group of recently diverged species where evolutionary signals are still detectable (Jenner and Wills 2007; Raff 2012). The nematode *Caenorhabditis elegans* was the first metazoan to have its genome sequenced, and consequently it is among the more thoroughly annotated and understood sequences available (Gerstein, et al. 2010). Recently, many genomes of close relatives of *C. elegans* have been sequenced, some of which have been assembled to chromosome-level contiguity (Fierst, et al. 2015; Kanzaki, et al. 2018; Ren et al. 2018; Yin, et al. 2018; Stevens, et al. 2019). Here, we harness these new resources to interrogate the evolution of repetitive genomic landscapes among five *Caenorhabditis* species. We find a conserved chromosomal structure of repetitive elements among four species, whereas the ecologically and morphologically divergent *C. inopinata* (Kanzaki, et al. 2018; Woodruff and Phillips 2018; Woodruff, et al. 2018) harbors an atypically uniform repetitive landscape driven by a handful of transposable element superfamilies.

## Methods

### Genome assemblies

Five *Caenorhabditis* genome assemblies with chromosome-level contiguity were used for this study (Fig. 1). The *C. elegans, C. briggsae*, and *C. nigoni* (Yin, et al. 2018) genomes were retrieved from WormBase ParaSite (parasite.wormbase.org; (Howe, et al. 2016)). The *C. inopinata* genome (Kanzaki, et al. 2018) was retrieved from the *Caenorhabditis* Genomes Project (Caenorhabditis.org; (Slos, et al. 2017; Stevens, et al. 2019)).

**Figure 1.**
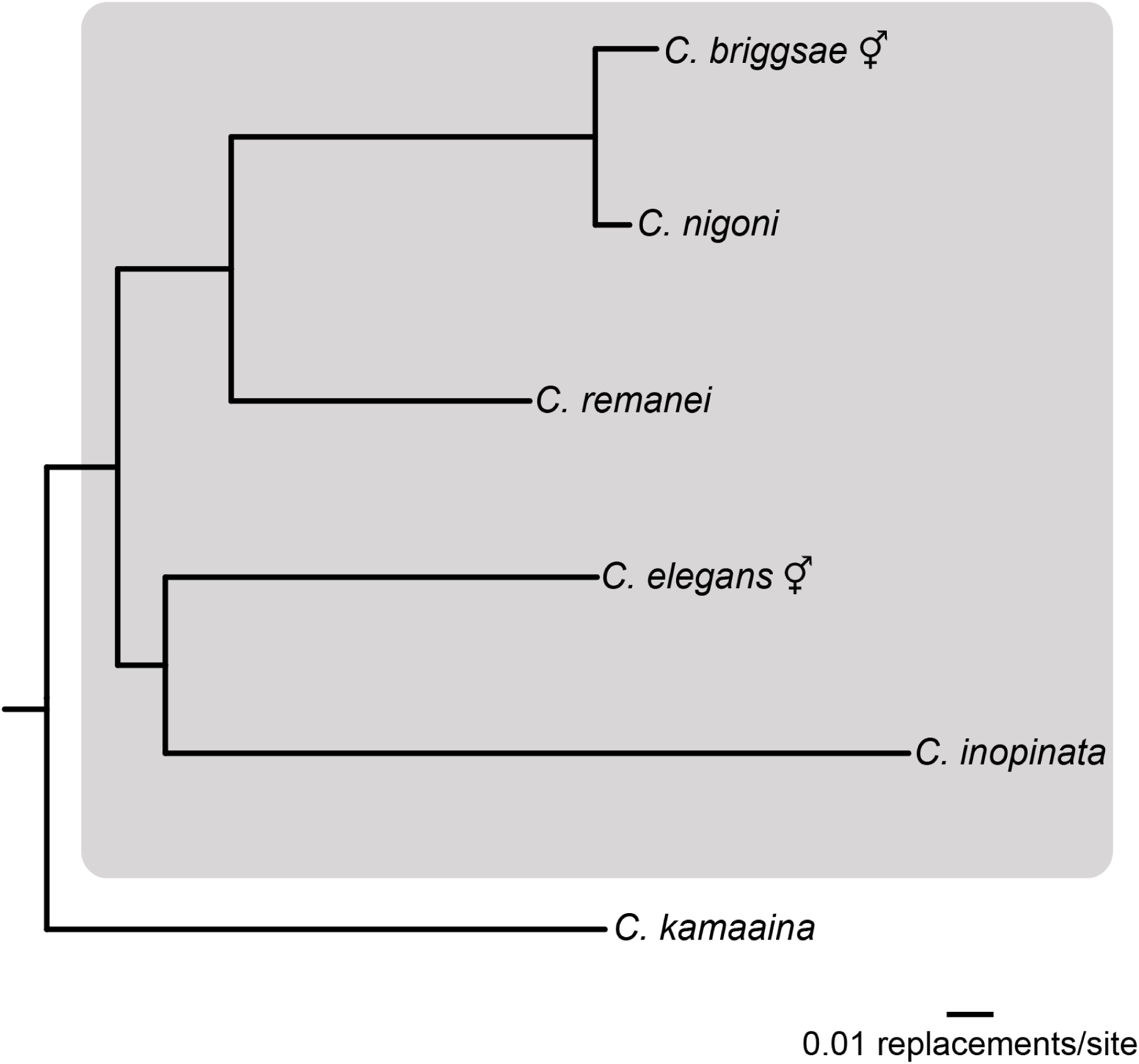
*Caenorhabditis* phylogeny. Species addressed in this study are included in the gray box. *C. elegans* and *C. briggsae* are hermaphroditic species (indicated by the symbol), whereas *C. nigoni, C. remanei*, and *C. inopinata* are female/male. The topology is derived from the Bayesian phylogeny inferred from protein sequences in (Stevens et al. 2019).

A new chromosome-level assembly of the *C. remanei* genome was kindly shared by Patrick Phillips and AAT (pers. comm.; genome manuscript in preparation). Protein sets of 28 *Caenorhabditis* species and *Diploscapter coronatus* were retreived from the *Caenorhabditis* Genomes Project (caenorhabditis.org). Versions of all genome assemblies and protein sets can be found with the deposited data and code associated with this work (https://github.com/gcwoodruff/transposable_elements_2019).

### Repeat inference and quantification

After excluding mitochondrial DNA, assemblies were masked and annotated for repeat content through a hybrid approach using multiple software packages (inspired by (Berriman, et al. 2018) and avrilomics.blogspot.com/2015/09/lrtharvest.html). De novo repeat libraries were generated with RepeatModeler (options-engine ncbi -pa 16; (Benson 1999; Bao and Eddy 2002; Price, et al. 2005)). Concurrently, sequences that align to transposable elements were detected with TransposonPSI using default parameters (transposonpsi.sourceforge.net). LTRHarvest was also used to identify LTR retrotransposon sequences in these assemblies (options -seqids yes -tabout no -gff3; (Ellinghaus, et al. 2008)). LTRHarvest output was further filtered (LTRHarvest option -hmms) to extract just those sequences containing only LTR retrotransposon domains; this was done with Pfam (Finn, et al. 2015) Hidden Markov Models of known LTR retrotransposon domains reported in (Steinbiss, et al. 2009). The RepeatModeler, TransposonPSI, and LTRHarvest species-specific repeat libraries were then concatenated with additional repeat libraries. One of these is the Rhabditida library (“Rhabditida.repeatmasker”) associated with the RepeatMasker software (Tarailo-Graovac and Chen 2009). Additionally, the *C. elegans* and *C. briggsae* repeat libraries (“cbrapp.ref,” “celapp.ref,” “cbrrep.ref,” and “celrep.ref”) from RepBase (Bao, et al. 2015) were also combined with the above libraries to generate redundant repeat libraries for each assembly. USEARCH (Edgar 2010) was then used to cluster repeats (options -id 0.8 -centroids –uc -consout -msaout) and generate non-redundant libraries. These repetitive sequences were then classified with RepeatClassifier (part of the RepeatModeler software) with default parameters. Repeats classified as “Unknown” were then aligned to *Caenorhabditis* species-specific protein sets with blastx (options -num_threads 24 -outfmt 6 -evalue 0.001; (Camacho, et al. 2009)); unclassified repeats that aligned to predicted proteins were then removed from the repeat libraries. These subsequent repeat libraries were then used with RepeatMasker (part of the RepeatModeler software; options -s -gff -pa 16) to mask the genome assemblies.

To measure and visualize the global landscape of repeats, Bedtools nuc (Quinlan 2014) was used to measure the fraction of masked bases in non-overlapping windows across the genomes (both 10 kb and 100kb window size; default parameters). This was also done for all specific transposable element classes, orders, superfamilies, and families. Repeats classified by RepeatClassifier (i.e., repeats not classified as “Unknown”) were annotated by a custom repeat taxonomy (repeat_taxonomy.tsv) informed by the classification system in (Wicker, et al. 2007). To measure the general trend of repeat density along all chromosomes simultaneously, windows were normalized by chromosome position using custom Linux scripts (all code associated with this work has been deposited (https://github.com/gcwoodruff/transposable_elements_2019)) by setting the median chromosome base pair to 0 and the end chromosome base pairs to 0.5. To further understand and quantify repetitive element genomic structure, chromosome arms were defined as genomic regions with normalized chromosome positions ≥ 0.25; chromosome centers were defined as regions with normalized chromosome positions < 0.25. To measure the impact of specific transposable element taxa on the global structure of repetitive elements, specific taxa were removed from the GFF file generated by RepeatMasker with custom Linux scripts, and assemblies were re-masked with Bedtools maskfasta with annotations excluding each specific family. Masked assemblies were then processed as above to quantify and visualize global repeat structure.

To understand the extent of transposon representation in protein-coding genes, all predicted protein sequences from 28 *Caenorhabditis* genomes were aligned to the TransposonPSI (transposonpsi.sourceforge.net) transposon protein database (“transposon_db.pep”) using blastp (options -outfmt 6 -evalue 0.001; (Camacho, et al. 2009)). Unique proteins that aligned to transposons were extracted and counted with custom Linux scripts to determine the fraction of transposon-aligning protein-coding genes per genome. In the case of *C. inopinata*, its transposon-aligning protein set was aligned to the set of *C. elegans* proteins (with its 100 transposon-aligning proteins removed) in the same manner to find the intersection of protein-coding genes that align to both transposons and to otherwise homologous nematode proteins.

Measures of the divergence of individual repetitive element insertions in *C. inopinata* were performed with phylogenetic analyses. Here, phylogenies were only performed on repetitive elements that had ≥100 insertions in the *C. inopinata* genome that were also not low complexity repeats. Repetitive elements of the same kind were defined as belonging to the same “cluster” as inferred by USEARCH above. Individual insertions were extracted from the repeat annotation files generated by Repeat-Masker with custom Linux scripts and Bedtools getfasta. Insertions derived from the same cluster were were aligned with MAFFT (options –auto and –adjustdirection) (Katoh and Standley 2013). Alignments were then trimmed with trimAl (option -automated1) (Capella-Gutiérrez, et al. 2009). Both trimmed and untrimmed alignments were then used to generate phylogenetic trees with FastTree (options -nt -gtr -quiet) (Price, et al. 2010). Before phylogenetic inference, trimmed alignments were filtered to exclude those <50 bp in length; this led to 265 clusters with untrimmed alignments being pared down to 165 clusters with trimmed alignments. The terminal branch lengths of all trees were extracted with the “phytools” R package (Revell 2012); the terminal branch length of each insertion was used as a measure of its divergence. Branch lengths were not converted to strict ages due to known dating issues in *Caenorhabditis* nematodes (Cutter 2008; Cutter, et al. 2019); these are likely exacerbated when using rapidly evolving transposable elements. All version information regarding the software used here can be found with the deposited data and code associated with this work (https://github.com/gcwoodruff/transposable_elements_2019).

### Simulations of transposable element evolution

Simulations of transposon element evolution were conducted in SLiM 3.3 (Haller and Messer 2018) with a script based on recipe 13.6 (“Modeling transposon elements”) from the SLiM 3.0 manual (28 February 2018 revision). In the recipe, active transposons that able to “copy and paste” themselves were simulated. Transposable elements are also disabled at some frequency, losing their ability to reproduce. We simulated the transposable element evolutionary dynamics with varying recombination landscapes and fitness effects depending on the genomic region of a transposon element insertion. For all simulations, the population size was 5,000 individuals. Genomes were modeled as a single 3 MB chromosome. Recombination was either uniform across the chromosome (r=1×10^−8^) or had three domains of different recombination rates in the chromosome arms and centers. In the case of three recombination domains, the chromosome arms had high a recombination rate (r=1×10^−8^) while chromosome centers had a lower recombination rate (r=1×10^−10^). The probability of transposable element replication/insertion was 1×10^−4^, and the probability of transposable element deactivation was 5×10^−5^. All simulations were run for 50,000 generations. All scenarios were replicated 50 times. From the simulations, we estimated the number of active and disabled transposable elements in the central and peripheral domains under different selection scenarios. All SLiM scripts for simulations have been deposited on Github (https://github.com/gcwoodruff/transposable_elements_2019). We studied the following scenarios:

a. All transposable element insertions are neutral (s = 0, Figure 8a);
b. Transposable elements are weakly deleterious; the fitness effects are drawn from a gamma distribution with mean s= −0.003 and alpha = 0.3 (Figure 8b);
c. Transposable elements are highly deleterious; the fitness effects are drawn from a gamma distribution with mean s= −0.03 and alpha = 0.3 (Figure 8c);
d. Transposable elements only located in the central domain are weakly deleterious; fitness effects are drawn from a gamma distribution with mean s= −0.003 and alpha = 0.3 (Figure 8d);
e. Transposable elements only located in the central domain are highly deleterious; fitness effects are drawn from a gamma distribution with mean s= −0.03 and alpha = 0.3 (Figure 8e);
f. Transposable elements located in the center are more deleterious than in the arms; fitness effects are drawn from a gamma distribution with mean s = −0.03 (centers) and mean s = −0.003 (arms) with alpha = 0.3 (Figure 8f);
g. Transposable elements located in the center are more deleterious than in the arms; fitness effects are drawn from a gamma distribution with mean s = equal −0.03 (centers) and mean s= −0.003 (arms) with alpha = 0.3 but TEs replicate via “cut-and-paste” (Figure 8g);
h. All transposable elements are weakly deleterious; fitness effects are drawn from a gamma distribution mean s= −0.003 and alpha = 0.3. In addition, highly beneficial mutations with fitness effects drawn from a gamma distribution with mean s = 0.1 and alpha = 0.3 occur with the mutation rate u = 1×10^−9^.

**Figure 2.**
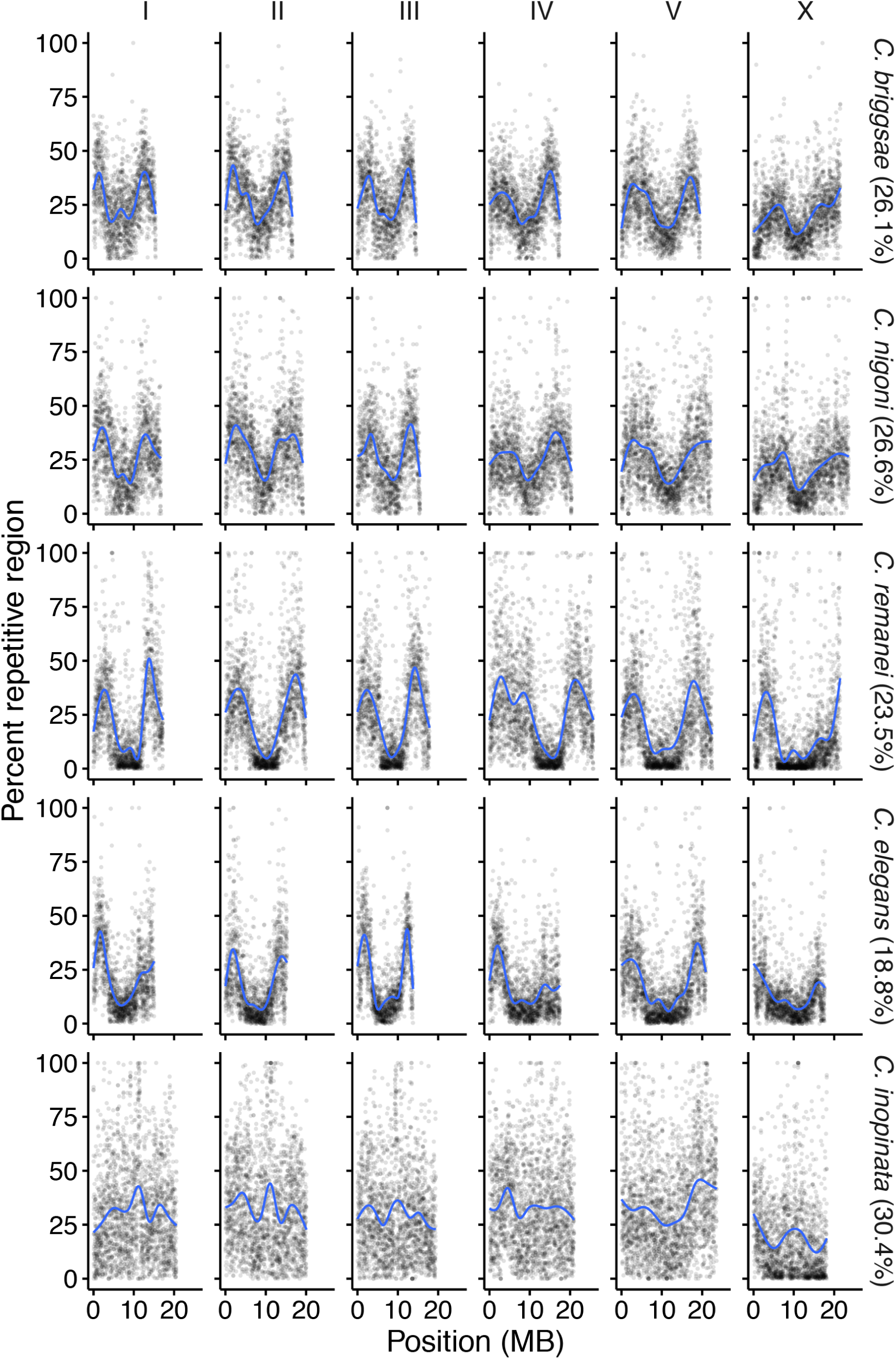
The genomic landscape of repetitive elements in five *Caenorhabditis* assemblies. Columns represent the six chromosomes; rows are the species ordered phylogenetically as in Figure 1. Plotted are the percentages of 10 kb windows that contain repetitive regions by genomic position. Percentage of the whole genome that is repetitive is noted in parentheses. Blue lines were fit with generalized additive models.

**Figure 3.**
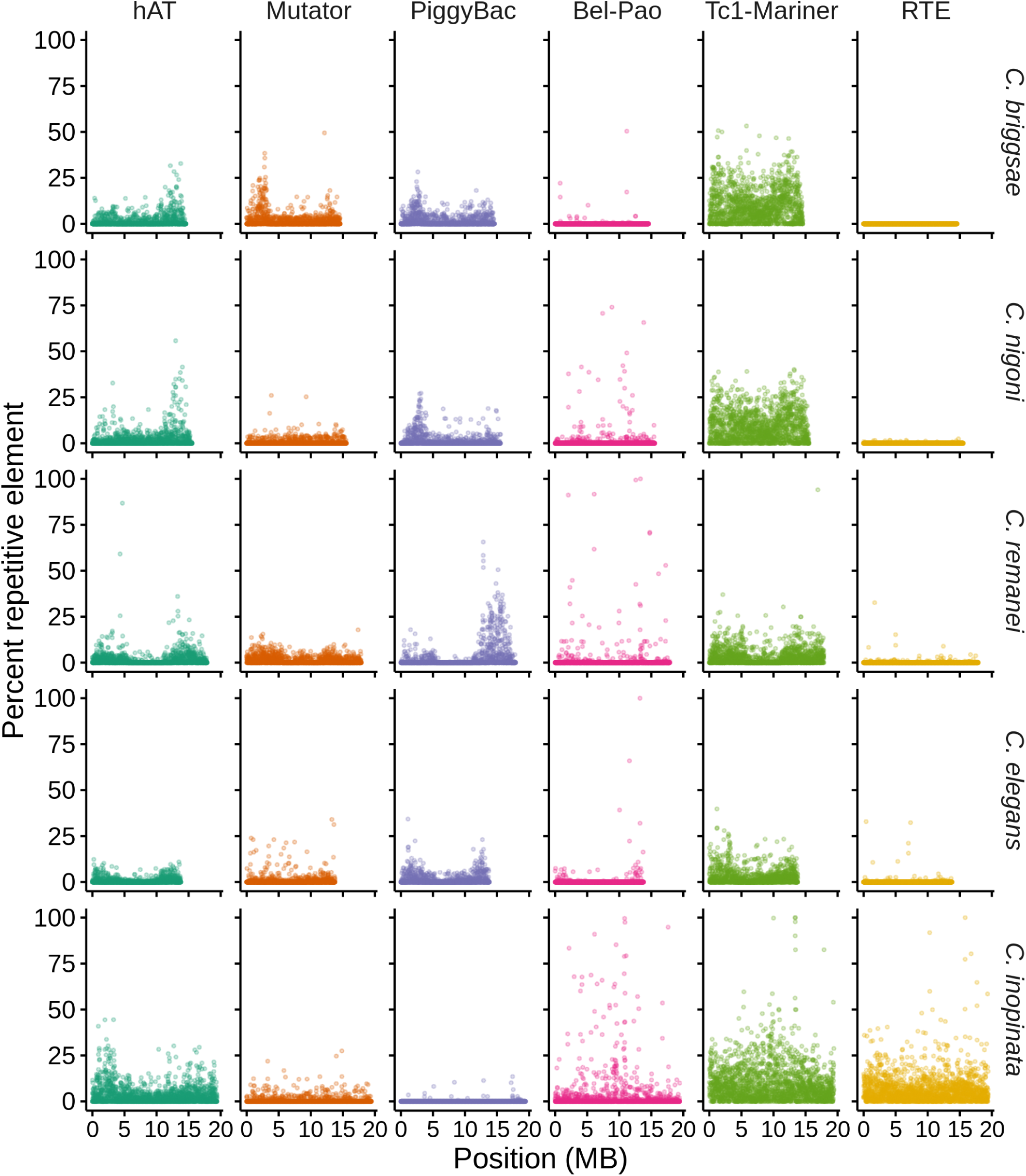
Repeat superfamilies vary in their chromosomal distributions. Plotted are the percentages of 10 kb windows that contain a given repeat superfamily (columns) along chromosome III in five species (rows). Six superfamilies among twenty-six detected were chosen to illustrate the diversity in repetitive landscapes (see Supplemental Material for all distributions of all repeat taxa on all chromosomes).

**Figure 4.**
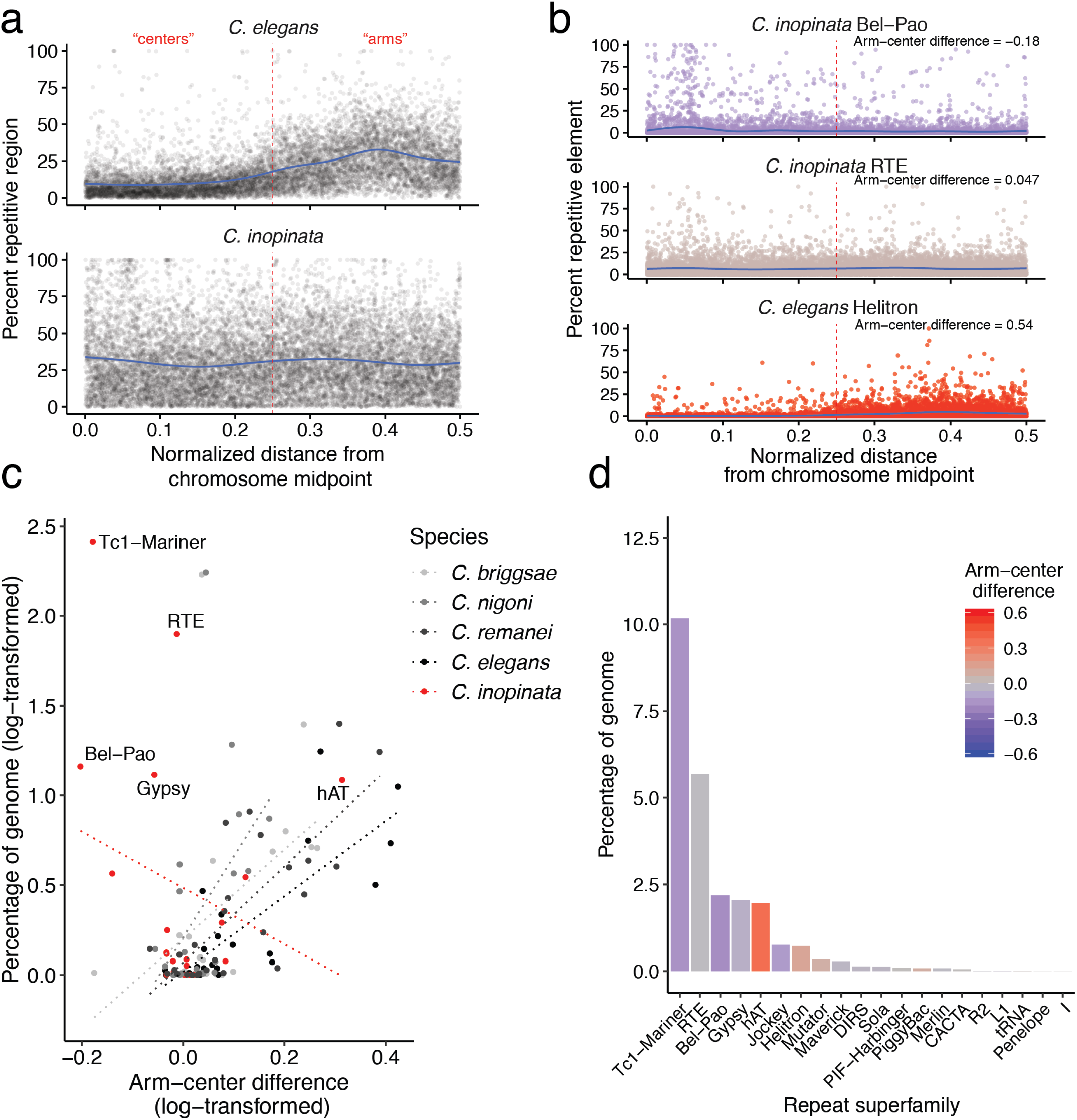
The genomic structure of repeat superfamilies in *Caenorhabditis*. a. The genomic landscape of repetitive elements in *C. elegans* and *C. inopinata* when normalized by chromosome position. Here, all genomic windows from all chromosomes are plotted, with the percentages of 10 kb windows that contain repetitive regions on the y-axis. “0” represents chromosome midpoints, and “0.5” represents chromosome ends. Windows can then be binned into chromosome “centers” (normalized chromosomal position < 0.25 (dotted red vertical line)) or “arms” (normalized chromosomal position greater than or equal to 0.25) to quantify the impact of chromosome position on repeat density. The blue lines were fit by generalized additive models. b. Quantifying the chromosomal structure of repeat taxa. The genomic distribution of three repeat superfamilies in among two species are plotted with normalized chromosomal positions as in (a) (from top to bottom: Bel-Pao in *C. inopinata*, RTE in *C. inopinata*, and Helitron in *C. elegans*). Points are colored by the Cohen’s *d* effect size of chromosome position (chromosome arms - centers) on repeat density with the same color gradient as in panel (d). Here and in all panels this is referred to as the “arm-center difference.” An effect size of 1 notes that the average repeat density among windows in chromosome arms is one pooled standard deviation higher than those in centers; an effect size of 0 reveals on average no difference in repeat density between chromosome arms and centers. Negative values reveal repeat densities higher in chromosome centers compared to arms. c. The relationship between repeat chromosomal structure and total genomic repeat content among repeat superfamilies in five *Caenorhabditis* species. The “arm-center difference” is the Cohen’s *d* effect size of chromosome position on repeat density as described in (b). The five most abundant repeat superfamilies in *C. inopinata* are labeled. All variables are log-transformed, (ln(variable+1)). All linear relationships p<0.05 except for *C. inopinata* (p=0.29); additional regression statistics can be found in the text and in the supplemental material. Supplemental Figure 11 shows the same data but not transformed. d. Repeat superfamily content in *C. inopinata*. Bars are colored by arm-center chromosome position effect size as described in panel (b).

**Figure 5.**
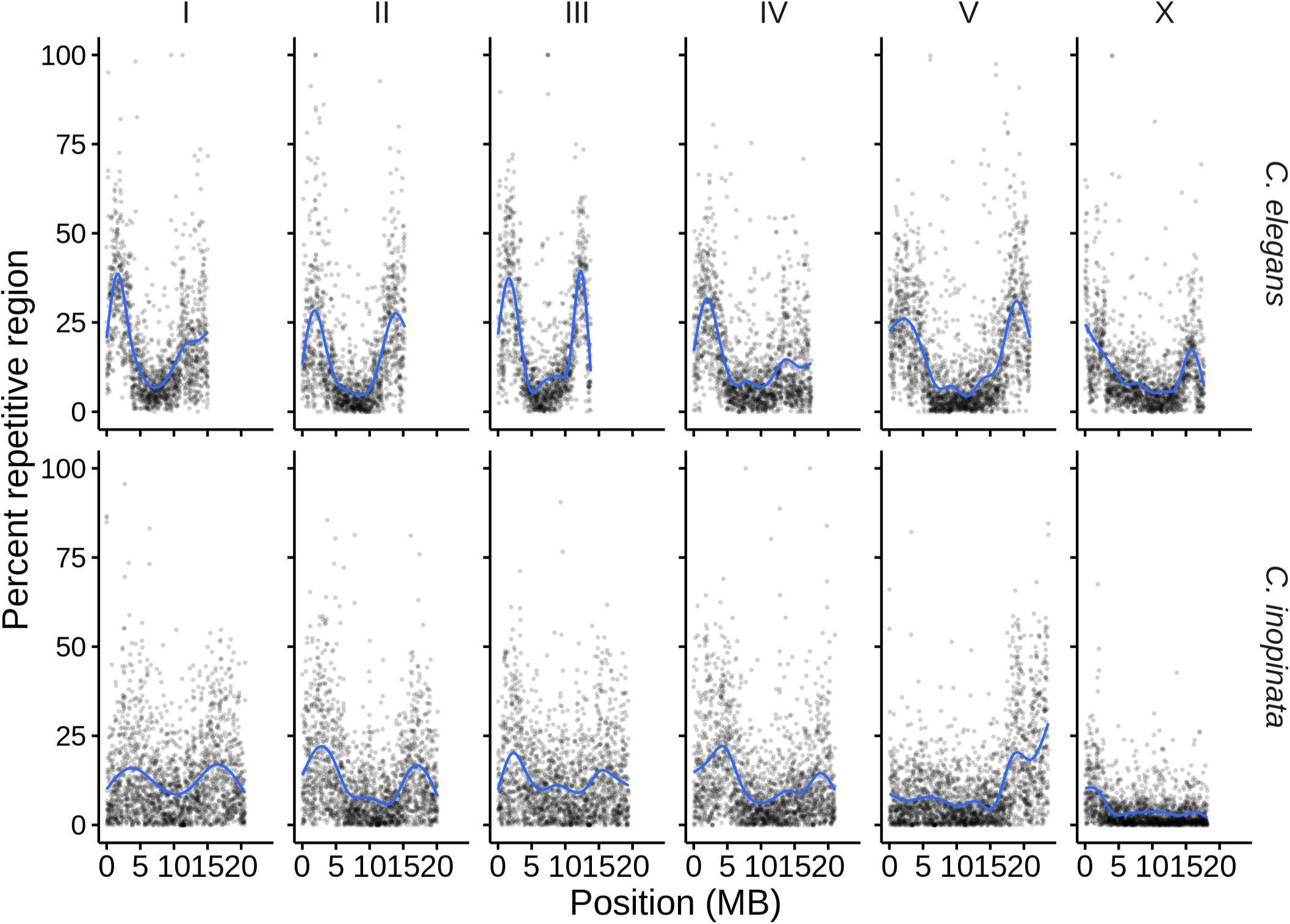
*C. inopinata* reveals a more conventional repetitive genomic landscape when four repeat superfamilies are removed. Plotted are the percentages of 10 kb windows that contain repetitive regions by genomic position after removing Tc1-Mariner, RTE, Bel-Pao, and Gypsy repeat superfamilies. The blue dashed line was fit by LOESS local regression.

**Figure 6.**
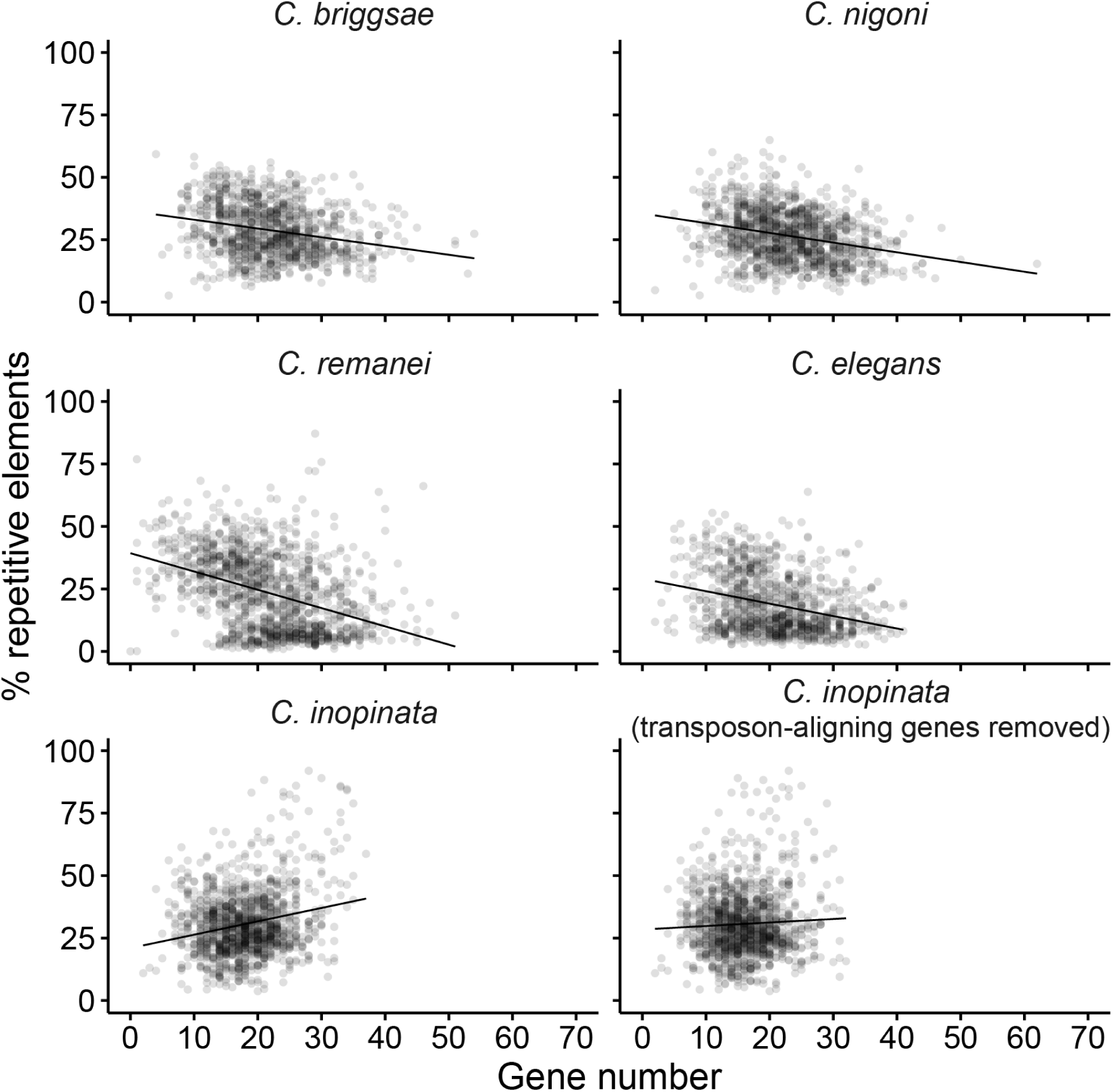
The relationship between gene and repeat density in *Caenorhabditis*. Plotted are the percent repetitive region by gene count in 100 kb windows across all species. In the case of *C. inopinata*, an additional plot excludes 2,489 transposon-aligning genes (that also do not align to any *C. elegans* proteins) from gene counts. All linear relationships p<0.0001 except for the last panel (p=0.070); additional regression statistics can be found in the text and in the supplemental material.

**Figure 7.**
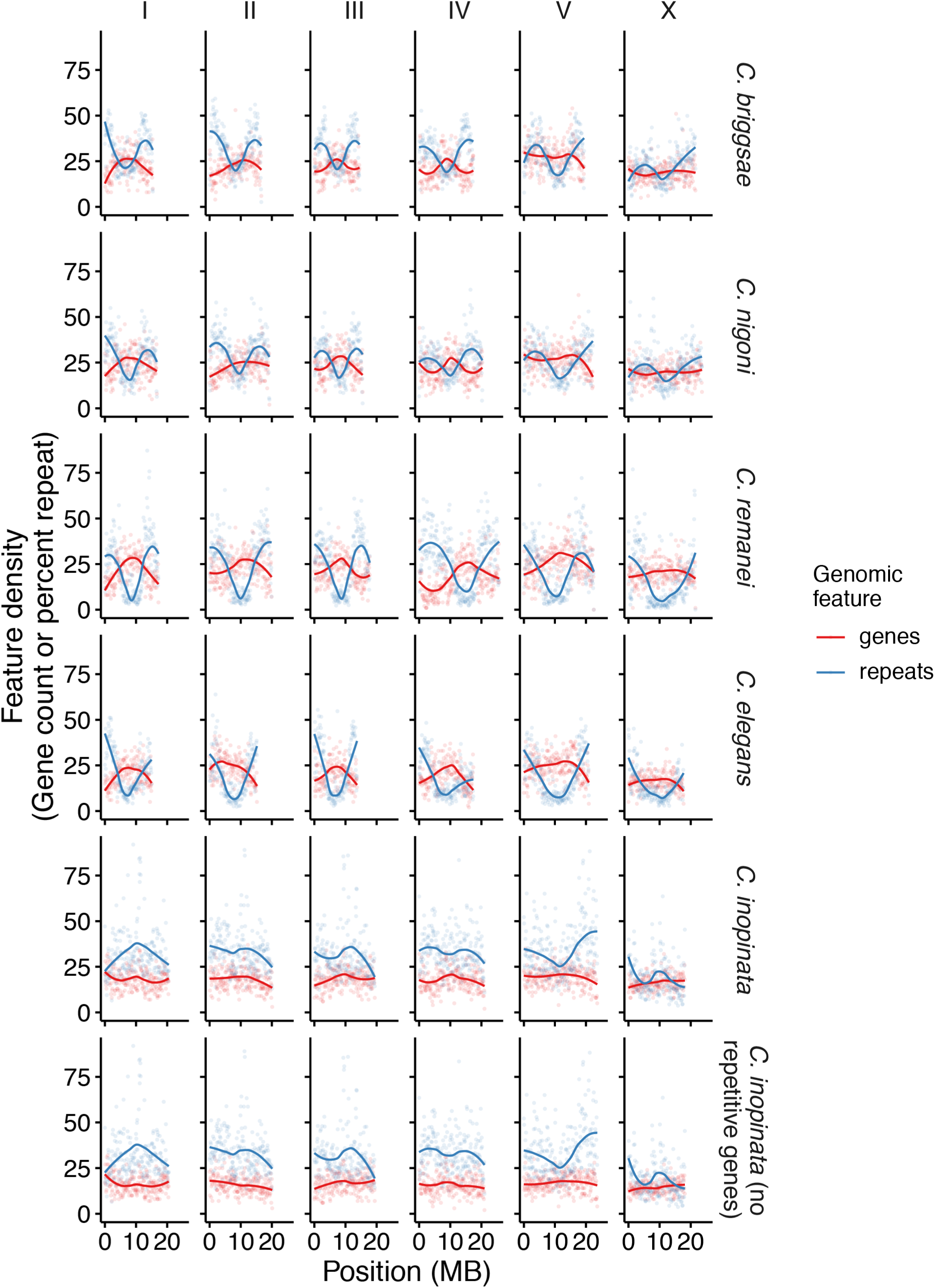
Genomic landscapes of gene and repeat density in *Caenorhabditis*. Columns represent the six chromosomes; rows represent species. Here, the gene densities of *C. inopinata* when 2,489 transposon-aligning genes are excluded are also plotted (last row; the repeat densities are identical to the row above). 100kb windows are plotted. For repeats, densities are reported as the percentage of the window that is repetitive. For genes, densities are reported as the number per window because gene length also covaries with genomic position (Supplemental Figures 21-22). Lines were fit by LOESS local regression.

**Figure 8.**
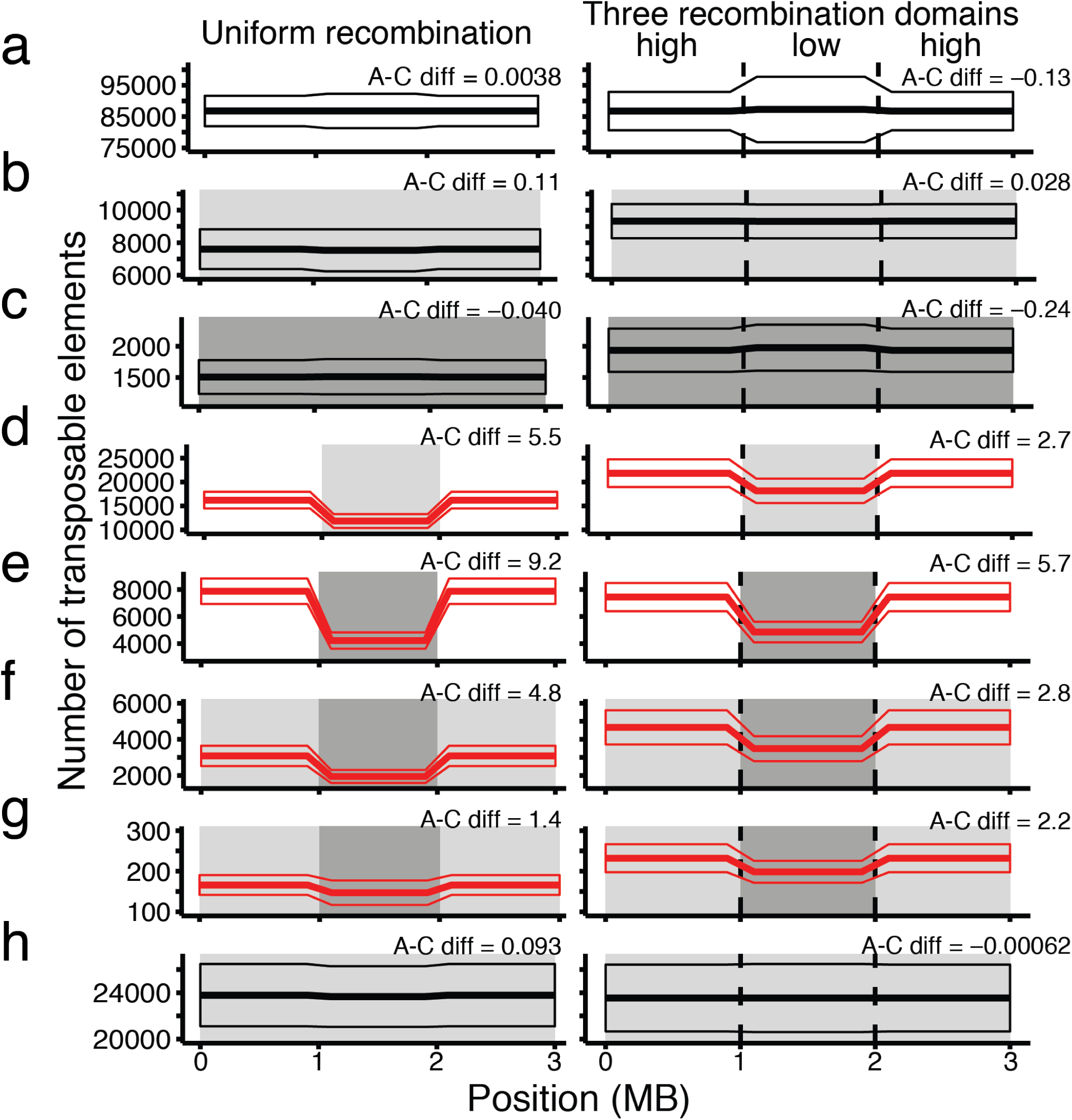
Simulations reveal chromosomal heterogeneity in insertion fitness effects is sufficient for promoting genomic repetitive organization. 3 MB chromosomes were evolved under multiple evolutionary scenarios. Plotted are the total number of transposable elements at four points along the chromosome after 50,000 generations. The left column shows scenarios where there were uniformly high rates of recombination along the chromosome (r= 1×10^−8^); the right column shows scenarios where there were three chromosomal domains of recombination (boundaries denoted by the dashed lines; chromosome arms r= 1×10^−8^; chromosome centers r= 1×10^−10^). Gray boxes represent chromosomal regions with deleterious fitness effects of transposable element insertion (light gray, mean s= −0.003; dark gray, mean s= −0.03; white, s= 0 for all transposable element insertions). When not neutral, fitness effects of insertion were drawn from a gamma distribution. Thick horizontal lines represent mean transposable element number among simulations (N simulations=50); thin lines represent 1 SDM. Red lines represent scenarios with significant arm-center Cohen’s d effect sizes (Wilcoxon rank-sum p=7.1×10^−18^-6.2×10^−9^); black lines show scenarios with no significant chromosomal organization of repeats (Wilcoxon rank-sum p=0.54-0.99). “A-C diff” is the arm-center Cohen’s *d* effect size of transposable element number between chromosome arms and centers. All scenarios implemented “copy-and-paste” models of transposable element replication with the exception of (g), which used a “cut-and-paste” model. Brief descriptions of each scenario are described below (see methods for details): a. All TE insertions neutral. b. All TE insertions weakly deleterious. c. All TE insertions highly deleterious. d. TE insertions weakly deleterious in center; TE insertions neutral in arms. e. TE insertions highly deleterious in center; TE insertions neutral in arms. f. TE insertions highly deleterious in center; TE insertions weakly deleterious in arms. g. TE insertions highly deleterious in center; TE insertions weakly deleterious in arms; “cut-and-paste” mode of TE replication. h. All TE insertions weakly deleterious; beneficial mutations introduced (u= 1×10^−9^).

### Statistical analyses

All statistical analyses and plots were generated with the R statistical language (Team 2019). The *lm* and *wilcox.test* functions in base R were used for linear models and performing Wilcoxon rank sum tests. The *cohen.d* function in the “effsize” (Torchiano 2019) package was used to estimate Cohen’s *d* effect sizes. The “ggplot2” (Wickham 2016), “lemon” (Edwards 2019), and “ggforce” (Pedersen 2019) R packages were used to make plots. Code and data for all analyses have been deposited in Github (https://github.com/gcwoodruff/transposable_elements_2019).

## Results

### Repeat density covaries with chromosomal position in all species but *C. inopinata*

The genomic landscape of repetitive elements was inferred in five *Caenorhabditis* assemblies (Fig. 1) through a combination of de novo and element class-specific methods (see methods; Fig. 2). For context, intrachromosomal heterogeneity of numerous genomic features has long been known in *C. elegans*, which has led to the definition of high-recombining, repeat-rich chromosome “arms” and low-recombining, gene-rich chromosome “centers” or “clusters” (Cutter, et al. 2009). These patterns appear to not be influenced by discrete centromeres as *C. elegans* has holocentric chromosomes where spindle attachment sites span the entire length of chromosomes (Albertson and Thomson 1982; Howe, et al. 2001). As previously described in nematodes (Duret, et al. 2000; Rizzon, et al. 2003; Stein, et al. 2003; Cutter, et al. 2009; Yin, et al. 2018), four of the assemblies reveal an enrichment of repetitive elements on chromosome arms relative to chromosome centers (Fig. 2; Supplemental Figure 1-2). Although there is a range in the repeat density difference between chromosome arms and centers among species (1-18% repeat content per 10kb window), *C. remanei* reveals the greatest difference in mean repeat content (Fig. 2; Supplemental Figure 1-2). Surprisingly, the closest relative of *C. elegans* used in this study, *C. inopinata*, revealed far less enrichment in chromosome arms (1% increased repeat content on arms compared to centers; Fig. 2; Supplemental Figure 1). After normalizing all genomic positions relative to chromosome centers, *C. inopinata* is the only species that did not reveal a significant relationship between chromosome position and repeat density in a linear model (Supplemental Figures 2-3; r^2^=1.8×10^−5^; *F*=1.2; β_1_= −1.5; p=0.27). Conversely, all other species had a positive relationship between repeat density and distance from chromosome center (Supplemental Figures 2-3; r^2^=0.078-0.24; *F*=999-3165; β_1_=32-60; p<0.0001 for all). There is then a largely conserved pattern of enrichment of repetitive elements in chromosome arms although it varies in its extent across species; *C. inopinata* is an exception in that it has almost no detectable global repeat chromosomal structure.

### Divergent genomic repetitive landscapes are driven by diversity in repeat taxon abundance and chromosomal structure

Transposable elements harbor abundant structural and replicative diversity. To understand the impact of such diversity on the genomic repetitive landscape, transposable elements were systematized into a taxonomy (repeat_taxonomy.tsv) informed by (Wicker, et al. 2007). The genomic landscapes of repeat taxa vary widely (Fig. 3; see Supplemental Material for the genomic distributions of all repeat taxa in all species). Repeat superfamilies (of which twenty-six were found among the repetitive elements in *Caenorhabditis*) can reveal conserved genomic landscapes with enrichment on chromosome arms (hAT superfamily, Fig. 3) or with little apparent chromosomal structure (Mutator superfamily, Fig. 3). Conversely, repeat superfamilies can vary widely in their genomic landscapes among species (such as PiggyBac, Bel-Pao, Tc1-Mariner, and RTE superfamilies; Fig. 2; Supplemental Material). Genomic repetitive landscapes are thus composed of dozens of repeat taxa that can each harbor idiosyncratic abundances and chromosomal structures among species and within genomes.

To further explore genomic landscapes of repeat taxa and their contributions to global repetitive structure, repeat taxon densities among chromosome arms and centers were compared (Fig. 4a-b; Supplemental Figures 4-10). Additionally, the total genomic abundances of various repeat taxa were also compared (Fig. 4c-d; Supplemental Figures 4-10; Supplemental Material). Among repeat classes (here including unclassified repeats, satellite DNA, and low complexity repetitive elements), transposable elements were most abundant (Supplemental Figure 4-5; Supplemental material), with class II DNA transposons generally being the dominant repeat class (Supplemental Figure 4-5). Consistent with global repeat structure, transposable elements tend to be enriched on chromosome arms (Supplemental Figure 4-5; Supplemental Material). However, in *C. inopinata*, class II DNA transposons are not enriched on chromosome arms (arm-center Cohen’s *d* effect size= 0.044, linear model p=0.69; Supplemental Figure 4-5), and class I DNA retrotransposons are enriched on chromosome centers, in exception to all other species (arm-center Cohen’s *d* effect size= −0.18, linear model p<0.0001; Supplemental Figure 4-5). Thus, variation in the chromosomal distributions and abundances of repeat classes underlies diversity in global repeat content.

Repeat taxa of lower ranks were interrogated further to understand the exceptional repetitive landscape of *C. inopinata*. All species but *C. inopinata* reveal a significant correlation between repeat superfamily genomic abundance and arm-center effect size (Figure 4c; Supplemental Figure 11; *C. inopinata*: r^2^=0.061, *F*=1.2, β_1_= −1.6, p=0.29;all other species: r^2^=0.21-0.64, *F*=5.0-38, β_1_=2.1-4.5, p=3.4×10^−6^-0.038). Some superfamilies that reveal atypical chromosomal structure are highly abundant in the *C. inopinata* genome, particularly Tc1-Mariner (class II transposon; 10.2% of the genome), RTE (class I LINE retrotransposon; 5.7%), Bel-Pao (class I LTR retrotransposon; 2.2%), and Gypsy (class I LTR retrotransposon; 2.0%; Fig. 4c-d), consistent with the previous description of the *C. inopinata* genome (Kanzaki, et al. 2018). Although these superfamilies are highly abundant and have atypical chromosomal structure, phylogenetic analyses suggest that insertions from these superfamilies are not on average any younger that those of other superfamilies in *C. inopinata* (Supplemental Figures 12-15). When these four superfamilies are excluded, the global repetitive landscape retains a chromosomal structure that resembles those of its close relatives (r^2^=0.048; *F*=621; β_1_=17.7; p<0.0001; Fig. 5). Thus, divergence in the repetitive genomic landscape can be driven in large part by the activity of a small number of repeat superfamilies that nonetheless differ greatly in structure and mode of replication.

### Gene density is negatively correlated with repeat content in all species but *C. inopinata*

Previous reports have noted that the genomic landscape of gene density mirrors that of repeat content in nematodes, and genes are enriched in chromosome centers relative to arms (Cutter, et al. 2009; Fierst, et al. 2015; Yin, et al. 2018). Non-random distributions of genes and repetitive elements also overlap with domains of high and low recombination (Rockman and Kruglyak 2009; Ross, et al. 2011). To explore if gene density is associated with repeat content in the assemblies addressed here, gene densities were estimated with publicly available genome annotations. When paired with estimates of repeat content, we found that, as expected, genes are moderately enriched in chromosome centers relative to arms in all five species (Supplemental figure 16-17; r^2^=0.025-0.12; *F*=33-180; β_1_= −6.2 to −0.20; p<0.0001 for all) and that gene density is negatively correlated with repeat content in four assemblies (Figure 6; r^2^=0.059-0.16; *F*=66-235; β_1_= −0.73 to −0.35; p<0.0001 for all but *C. inopinata*). However, *C. inopinata* revealed a positive correlation among repeat content and gene density (Figure 6; r^2^=0.050; *F*=65; β_1_=0.54; p<0.0001). As the previous description of the *C. inopinata* genome mentioned a transposable element insertion within the highly-conserved sex determination protein-coding gene *her-1* (Kanzaki, et al. 2018), we suspected this positive relationship may be due to the presence of repetitive content in the predicted protein-coding gene set. To test this, we aligned all predicted protein-coding genes from 28 Caenorahbditis genomes and one outgroup (*Diploscapter coronatus* (Hiraki, et al. 2017)) to the TransposonPSI transposon protein database. In most *Caenorhabditis* genomes, only a small fraction of predicted protein-coding genes aligns to transposon-related proteins (median= 1.08%; IQR=0.47%). However, in *C. inopinata* and *C. japonica*, a substantial fraction of their protein sets aligns to such proteins (Supplemental Figure 18; *C. inopinata*, 15%; *C. japonica*, 12%). Within *C. inopinata*, of the 3,349 proteins that aligned to the transposon database, 860 also aligned to *C. elegans* non-repetitive proteins. Indeed, after *C. inopinata* proteins aligning exclusively to transposable elements are removed, there is no significant relationship between repeat content and gene density (Figure 6; r^2^=0.002; *F*=3.3; β_1_=0.14; p=0.070). Moreover, this also abolishes the relationship between gene content and chromosomal position in this species (Figure 7; Supplemental Figures 16-17, 19-20; r^2^= −0.00090; *F*=0.89; β_1_= - 0.90; p=0.35). In tandem with patterns of repeat content, this is suggestive of a radical remodeling of genomic structure along the *C. inopinata* lineage.

### Simulations reveal chromosomal heterogeneity in insertion fitness effects is sufficient for promoting non-random genomic repetitive organization

Recombination rate also covaries with chromosomal position in nematodes and is elevated in chromosome arms relative to centers in both *C. elegans* and *C. briggsae* (Rockman and Kruglyak 2009; Ross, et al. 2011). How does intrachromosomal variation in recombination rate contribute to genomic repetitive organization? To address this question, we implemented forward simulations of transposable element evolution with the SLiM software package (Haller and Messer 2018; Figure 8; Supplemental material; see methods). Primarily, chromosomes with either a uniformly high recombination rate or with three domains of differing recombination rate (high in chromosome arms and low in centers) were modeled. In addition, various patterns of chromosomal heterogeneity in the fitness effects of transposable element insertion were also addressed. Notably, intrachromosomal variation in recombination alone was insufficient to generate chromosomal heterogeneity in transposon density. This was true in all simulations where there was no within-chromosome variation in the fitness effects of transposable element insertion (Figure 8a-c, h; Arm-center Cohen’s *d* effect size= −0.24-0.11; Wilcoxon rank sum test W=1056-1338, *p*= 0.54-0.99). Conversely, heterogeneity in insertion fitness effects consistently revealed genomic organizations of repetitive elements reminiscent of those observed in nematodes (Figure 8d-g; Arm-center Cohen’s *d* effect size=1.4-9.2; Wilcoxon rank sum test W=1803-2500, *p*= 7.1×10^−18^-6.2×10^−9^). This was observed even in the absence of chromosomal domains of differing recombination rates (Figure 8d-g, left column; Arm-center Cohen’s *d* effect size=1.4-9.2; Wilcoxon rank sum test W=1803-2500, *p*=7.1×10^−18^-6.2×10^−9^). Specifically, repeats are enriched in chromosomal regions where insertions are more deleterious than other chromosomal regions. This holds for both “copy-and-paste” as well as “cut-and-paste” (Figure 8g; Arm-center Cohen’s *d* effect size=1.4, 2.2; Wilcoxon rank sum test W=2139, 2368, *p*=9.1×10^−9^, 1.3×10^−14^) models of transposable element replication. Furthermore, the introduction of beneficial mutations is also insufficient to generate heterogenic genomic organizations of transposable elements (Figure 8h; Arm-center Cohen’s *d* effect size= −0.00062, 0.093; Wilcoxon rank sum test W=1247, 1293, *p*=0.99, 0.77). As transposable element insertion can abrogate gene function, the enrichment of gene density in chromosome centers is then a potential contributor to genomic repetitive organization.

## Discussion

Mobile elements are known to harbor tremendous diversity in their replication strategies. However, it remains unclear to what extent repeat taxon-specific evolution shapes the abundance and chromosomal distribution of global repetitive content among genomes. Here, we described just these patterns of transposable element content in five *Caenorhabditis* assemblies, and repeat taxon-specific evolution (repeat_taxonomy.tsv) among lineages appears widespread in this group. Diversity in the abundances of various repeat superfamilies is common. For instance, the Jockey TE superfamily has expanded in the *C. remanei* lineage; the Helitron superfamily of DNA transposons is uniquely abundant in the *C. briggsae*/*C. nigoni* clade (Supplemental Figure 8; Supplemental material). There is broad variation in repeat content both among and within genomes across the range of TE taxa (Fig. 3; Supplemental Figures 4-10; Supplemental material). Such chromosomal heterogeneity in repeat density depending on the type of transposable element has also been observed in multiple systems, including humans (Myers, et al. 2005), mice (Shifman, et al. 2006), and maize (Stitzer, et al. 2019). However, here elements generally tend to be enriched on chromosome arms relative to centers, as has observed in numerous studies in *Caenorhabditis* (Figure 2; Fig. 4c; Supplemental Figure 11; Supplemental material; (Consortium 1998; Stein, et al. 2003; Fierst, et al. 2015; Yin, et al. 2018)).

*C. inopinata* obviously bucks this general trend in repeat organization. Although its high repeat content has been previously noted (Kanzaki et al. 2018), here we describe its exceptional repetitive landscape. Despite this divergent genomic structure, some repeat taxa in this species exhibit the conventional enrichment on chromosome arms, primarily hAT, Mutator, and Helitron DNA transposons (Fig. 4d; Supplemental Figure 6; Supplemental material). Additionally, repetitive regions not obviously related to transposable elements (such as Satellite DNA, low complexity repeats, and unclassified repetitive elements) are also enriched on chromosome arms in *C. inopinata* (Supplemental Figure 4; Supplemental material). Thus the breakdown in the repetitive genomic landscape in this species is repeat taxon-specific. The exclusion of four repeat superfamilies that have high abundance and atypical genomic organization reveal a more conserved repeat structure (Fig 4c-d; Fig. 5). Notably, these particular repeat superfamilies vary greatly in their replicative diversity. Tc1/Mariner elements are cut-and-paste class II DNA transposons (Wicker, et al. 2007). RTE is a class I LINE retrotransposon that uses its own transcription product to prime reverse transcription (Lynch 2007; Wicker, et al. 2007). Bel-Pao and Gypsy are more complex LTR retrotransposons whose autonomous copies usually encode retroviral-derived proteins and use tRNA’s to prime reverse transcription (Lynch 2007; Wicker, et al. 2007). Thus variation in the abundance and chromosomal distribution of diverse repeat taxa underlies change in repetitive landscapes.

This chromosomal heterogeneity in transposable element distribution among nematode chromosomes has been widely noted (Duret, et al. 2000; Rizzon, et al. 2003; Cutter, et al. 2009), and these patterns covary with multiple genomic features that are also non-randomly distributed along chromosomes (Cutter, et al. 2009). In *C. elegans*, recombination rate is elevated on chromosome arms (Rockman and Kruglyak 2009), and protein-coding genes are enriched on chromosome centers (Consortium 1998). Heterochromatic regions (Garrigues, et al. 2015) and piRNA’s (Shi, et al. 2013) are also enriched on chromosome arms; consistent with this, chromosome arms are less transcriptionally active than chromosome centers (Garrigues, et al. 2015). As heterochromatin and small RNA’s have been connected to transposon regulation in other systems (Lee 2015), transposable elements may proliferate in such genomic regions where insertions are less costly and easier to control. Further, essential genes are enriched in chromosome centers in *C. elegans* (Kamath, et al. 2003). And, chromosome arms are associated with each other in physical space and tend to be tethered to nuclear membranes (Cabianca, et al. 2019). Such patterns of these genomic features have been observed across *Caenorhabditis* (Stein, et al. 2003; Ross, et al. 2011; Fierst, et al. 2015; Kanzaki, et al. 2018; Yin, et al. 2018). Taken together, this suggests that this chromosomal organization is an ancient, defining characteristic of the *Caenorhabditis* genome. This may also explain the counterintuitive observation that repetitive elements are enriched in high recombining regions in this system, whereas opposite patterns are observed in other systems (Rizzon, et al. 2002; Wright, et al. 2003; Jensen-Seaman, et al. 2004; Liu, et al. 2009; Pan, et al. 2011) and are predicted by evolutionary theory (Hill and Robertson 1966; Langley, et al. 1988). Although it is difficult to disentangle cause and effect, the heterochromatic, less gene-dense chromosomal regions have higher rates of recombination; repetitive elements accumulate there perhaps because of their potentially lower fitness effects upon insertion.

We performed evolutionary simulations to understand how heterogeneity in recombination rate and insertion fitness effects along the chromosome can influence repetitive genomic landscapes (Figure 8). Notably, we found that chromosomal heterogeneity in recombination rate alone is insufficient for promoting heterogeneity in transposable element content (Fig. 8a-c, h). This is largely consistent with previous evolutionary simulations that suggested that extremely low recombination rates are needed for Hill-Robertson effects to fix transposable elements (Dolgin and Charlesworth 2008). Instead, chromosomal domains of differing fitness effects of insertion were sufficient for promoting such patterns of repetitive content (Fig. 8d-g). This suggests that stronger purifying selection on transposable element insertions in the genomic clusters of genes in the centers of chromosomes are driving the conserved repetitive genomic landscapes in *Caenorhabditis*. Indeed, *C. inopinata* does not have protein-coding genes enriched in the centers of chromosomes like its close relatives (Fig. 7; Supplemental Figures 16-19, 19-20), which is consistent with its divergent repetitive genomic landscape and this hypothesis. However, not all transposable element taxa have lost genomic structure in *C. inopinata* (Fig. 3), so the weakened chromosomal heterogeneity in protein-coding gene content cannot explain its repetitive genomic landscape entirely. It is possible that the four exceptional repeat superfamilies expanded recently, after *C. inopinata* lost its central chromosomal clusters of protein-coding genes. However, phylogenetic approaches provide no evidence that such superfamilies are on the whole much younger than other transposable element taxa in this species (although there are a number of young insertions, Supplemental Figures 12-15). The role of small RNA regulators on the evolution of *Caenorhabditis* repetitive landscapes, and these types of elements in particular, is also an important open question because a number of such genes have been lost in *C. inopinata* (Kanzaki, et al. 2018). Particularly, three genes in the ERGO-1 siRNA pathway (*ergo-1, eri-6*/*eri-7*, and *eri-9*; (Piatek and Werner 2014)) are not present in this species (Kanzaki, et al. 2018). As *ergo-1* encodes an RNA-silencing argonaut protein (Yigit, et al. 2006), and as small RNA’s are known to regulate transposable elements in other systems (Piatek and Werner 2014), the loss of these genes in *C. inopinata* poses a tantalizing line of future research for understanding its repetitive landscape. Furthermore, the demographic and evolutionary history of a lineage can drive variation in the distributions of transposable elements (Lockton, et al. 2008), and this remains unexplored in *C. inopinata*. In any case, future work on recombination and the genomic organization of various features in *C. inopinata* (such as heterochromatin, spatial genomic structure, and transcription, among others) within their evolutionary context will be needed to better understand the causes of its divergent repetitive genomic landscape.

*C. inopinata* was not a lone outlier in one respect– *C. japonica* was also observed to have a high percentage of its gene set aligning to transposable elements (12%; Supplemental Figure 18). Previously observed to have high transposable element content (Fierst, et al. 2015; Szitenberg, et al. 2016), *C. japonica* shares notable ecological features with *C. inopinata*. They have only been observed on east Asian islands—*C. japonica* on Kyushu Island in Japan (Kiontke, et al. 2002; Yoshiga, et al. 2013) and *C. inopinata* in Okinawa and Taiwan (Kanzaki, et al. 2018; Woodruff and Phillips 2018). Moreover, they are both phoretic host specialists. *Caenorhabditis* nematodes generally thrive on rotting plant material (Kiontke, et al. 2011), and they travel from one resource patch to another on invertebrate carriers (Schulenburg and Félix 2017). Many *Caenorhabditis* species are promiscuous with respect to their hosts (including *C. elegans, C. briggsae, C. remanei*)(Cutter 2015), whereas others appear to be host specialists, having only been found with one invertebrate carrier. *C. japonica* is associated with the Japanese shield bug *Parastrachia japonensis* (Okumura, et al. 2013; Yoshiga, et al. 2013), and *C. inopinata* disperses on *Ceratosolen* fig wasps (Kanzaki, et al. 2018; Woodruff and Phillips 2018). The proliferation of transposable elements in this group may then somehow connected to host specialization. However, some presumptive host specialists do not have obvious expansion of transposable element content among their predicted protein-coding genes (*C. angaria, C. plicata*, and *C. monodelphis*; Supplemental Figure 18; Volk 1951; Sudhaus, et al. 2011; Slos, et al. 2017), and the extent of host preferences among *Caenorhabditis* species remains largely unknown (Cutter 2015). Additionally, there are many potentially island-restricted *Caenorhabditis* species whose genomes have yet to be assembled (Cutter 2015). Thus, the generation of more genome assemblies, in tandem with further exploration of *Caenorhabditis* ecology, will be needed to frame this question in its proper comparative phylogenetic context and address the possible role of ecological specialization in the evolution of transposable elements.

Such detailed phylogenetic comparative methods using more taxa would also be needed to address the obvious question regarding the impact of reproductive mode on transposable element content (Glémin, et al. 2019). Evolutionary theory predicts conflicting expectations regarding the abundance of transposable elements in selfing lineages (Wright and Schoen 1999; Morgan 2001), with different models of selection leading to either transposon expansion or reduction. Additionally, selfing lineages of *Arabidopsis* harbor more transposable elements than outcrossers (Wright, et al. 2001; Lockton and Gaut 2010). But in agreement with previous results (Fierst, et al. 2015; Yin, et al. 2018), we find no obvious pattern among transposable element content and reproductive mode, although our phylogenetic sample is quite small. This is also consistent with observations in asexual arthropods (Bast, et al. 2015) and dbelloid rotifers (Nowell, et al. 2018), that reveal no amplification of repeat content upon change in reproductive mode. As the *Caenorhabditis* genus only has three independent transitions to self-fertile hermaphroditism (Stevens, et al. 2019), this group may not be well-suited for addressing this question in a comparative phylogenetic context. However, one study reported that transposable element content among a broad phylogenetic sample of nematodes found no clear predictors of transposable element abundance and concluded that genetic drift is a major driver of transposon evolution (Szitenberg, et al. 2016). Although we also see no clear association of transposable element abundance with predicted population sizes here, as they range across at least two orders of magnitude in the *Caenorhabditis* genus (Dey, et al. 2013; Fierst, et al. 2015), future comparative phylogenetic approaches can also be used to test the prediction that genetic drift should influence transposable element abundance.

Regardless of the forces underlying their proliferation, the exceptional transposable element expansion in the exceptionally-morphological *C. inopinata* lineage cannot be ignored. Whereas many *Caenorhabditis* species are largely morphologically indistinguishable, *C. inopinata* is twice as long and develops twice as slowly as its close relatives (Kanzaki, et al. 2018; Woodruff, et al. 2018; Woodruff, et al. 2019), and it thrives in the lumen of fresh figs instead of on rotting plant material (Kanzaki, et al. 2018; Woodruff and Phillips 2018). Its divergent genomic structure is likewise striking. Repetitive elements have been shown to be a source of adaptive mutations in numerous contexts—peppered moths (van’t Hof, et al. 2016), oranges (Butelli, et al. 2012), and stickleback fish (Ishikawa, et al. 2019) all have evidence of such beneficial insertions. In addition to laying the groundwork for understanding the origins, maintenance, and stability of genome structure in the face of rampant mobile element proliferation, this work also sets the stage for understanding how mobile elements can promote or constrain rapid morphological and ecological change.

## Supporting information

Supplemental tables and statistics

## Supplemental Material

**Supplemental Figure 1.**
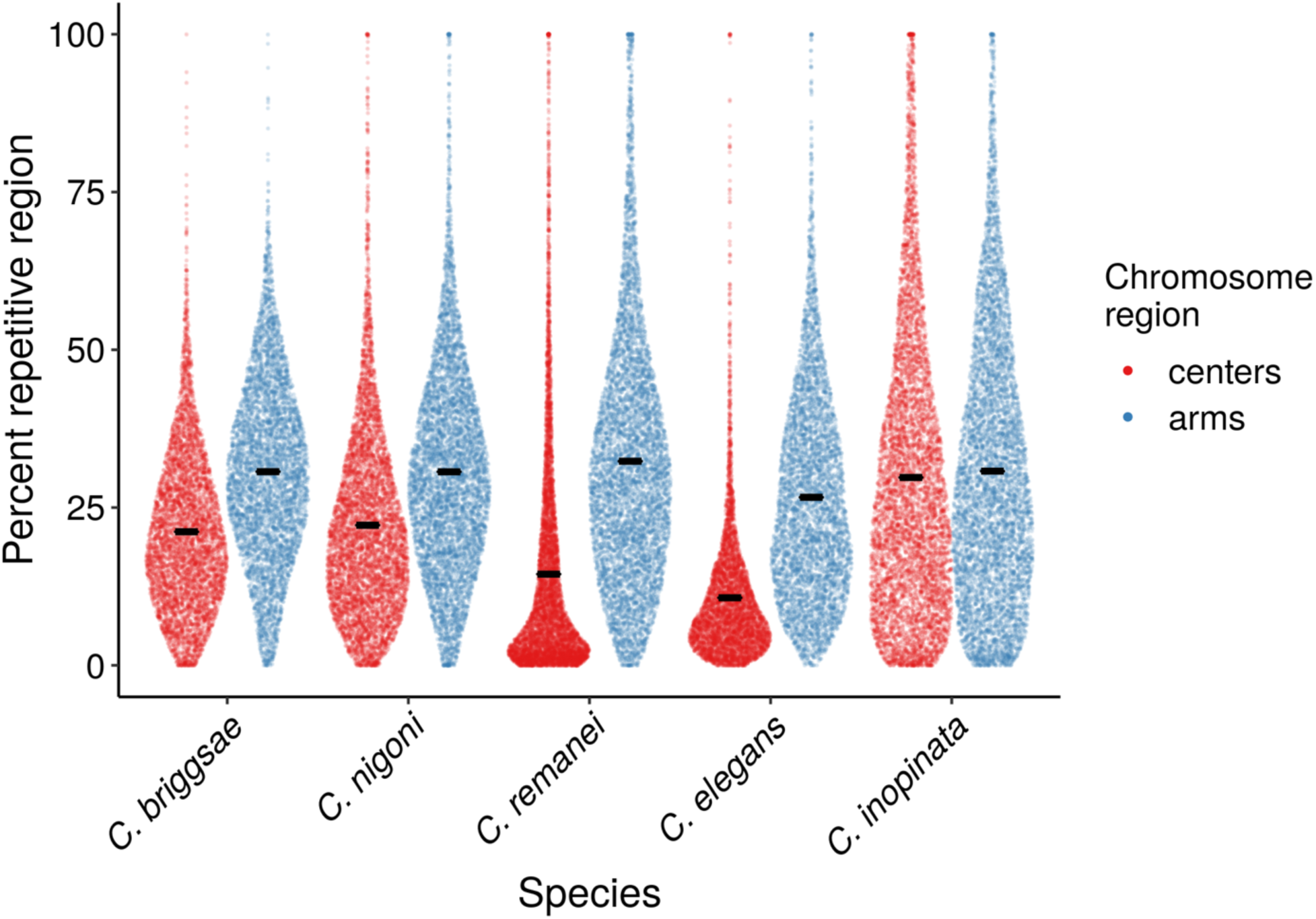
Distribution of repeat densities on chromosome arms and centers. Each point represents one 10 kb genomic window. After normalizing chromosome positions by distance to chromosome midpoint, windows were classified as being in chromosome “centers” (middle half of chromosome) or “arms” (outer half of chromosome). Sina plots (strip charts with points taking the contours of a violin plot) reveal the distribution of repeat densities of these windows in chromosome centers and arms for all species. Black horizontal lines, means.

**Supplemental Figure 2.**
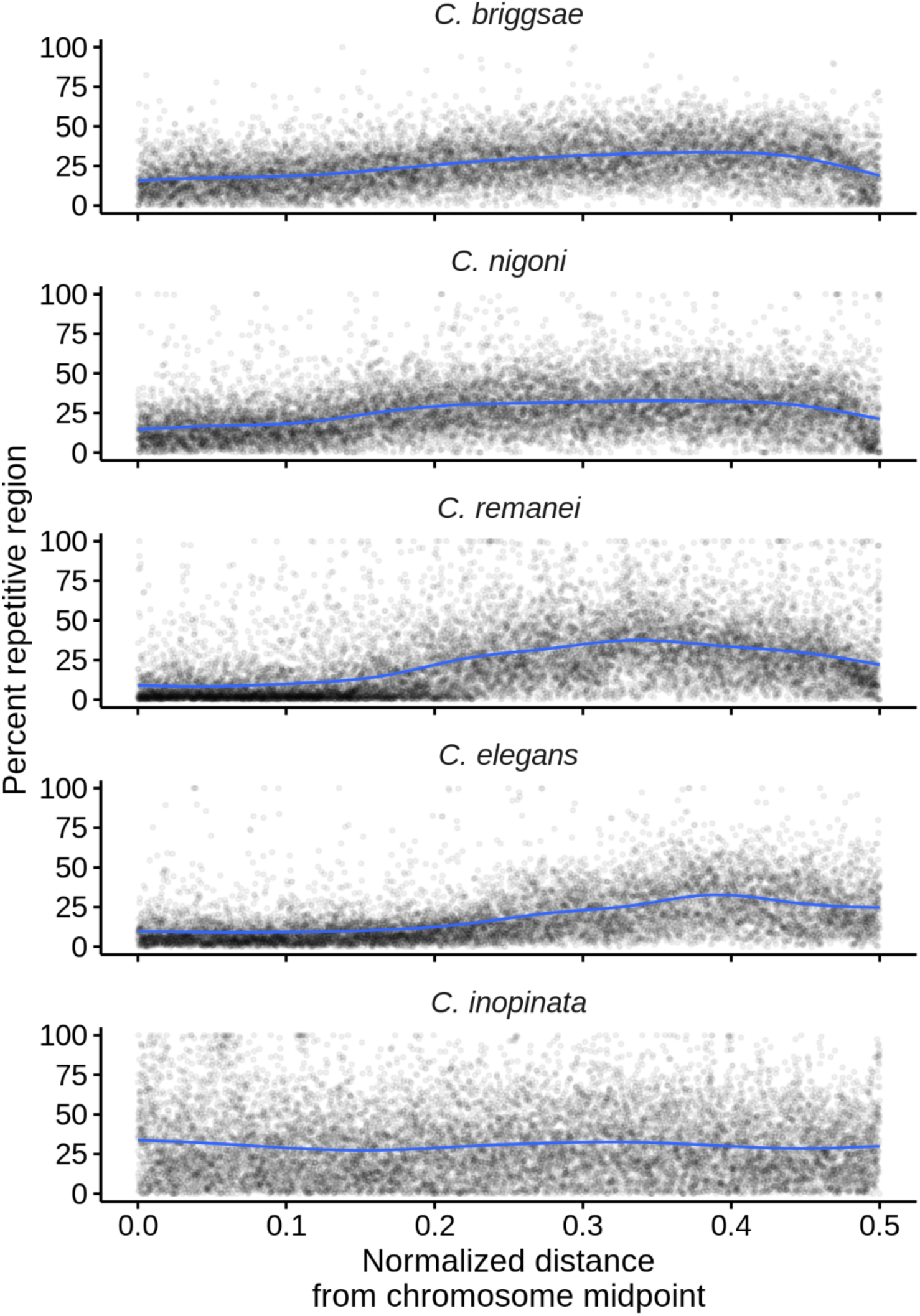
The genomic landscape of repetitive elements in *Caenorhabditis* after normalizing genomic position by distance from chromosome midpoints. Here, all chromosomes are plotted, and “0” on the x-axis represents chromosome midpoints. Columns represent the six chromosomes; rows are the species ordered phylogenetically as in Figure 1. Plotted are the percentages of 10 kb windows that contain repetitive regions by genomic position. Blue lines are fit by generalized additive models.

**Supplemental Figure 3.**
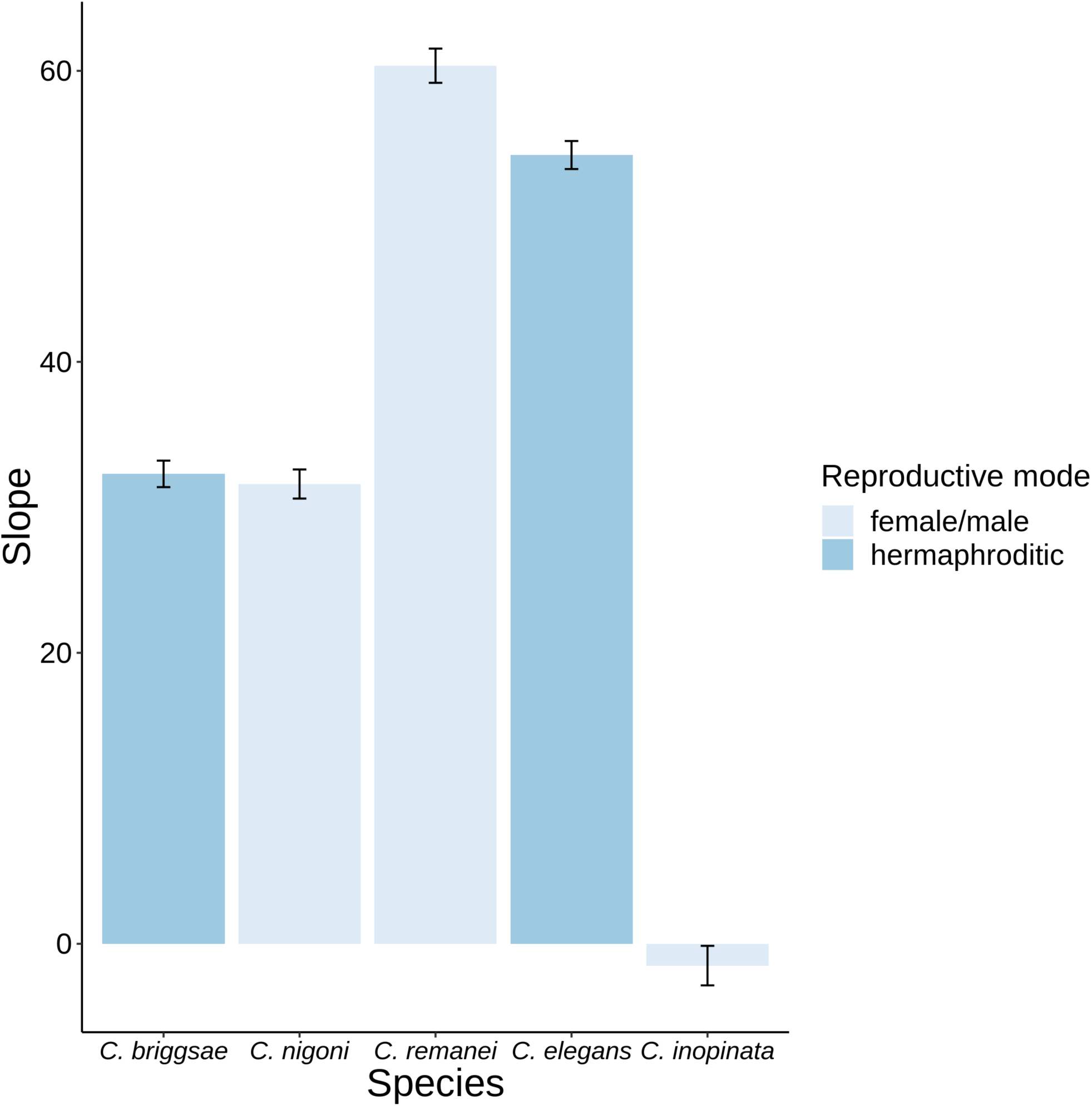
Coefficients (slopes) of linear models of the relationship between repeat density (percentage of window repetitive) and normalized distance from chromosome center (range=0-0.5; “0” is the chromosomal midpoint). All model *p*<0.0001 except for *C. inopinata* (*p*=0.27). See Supplemental Material (supplemental_statistics.xls) for model statistics.

**Supplemental Figure 4.**
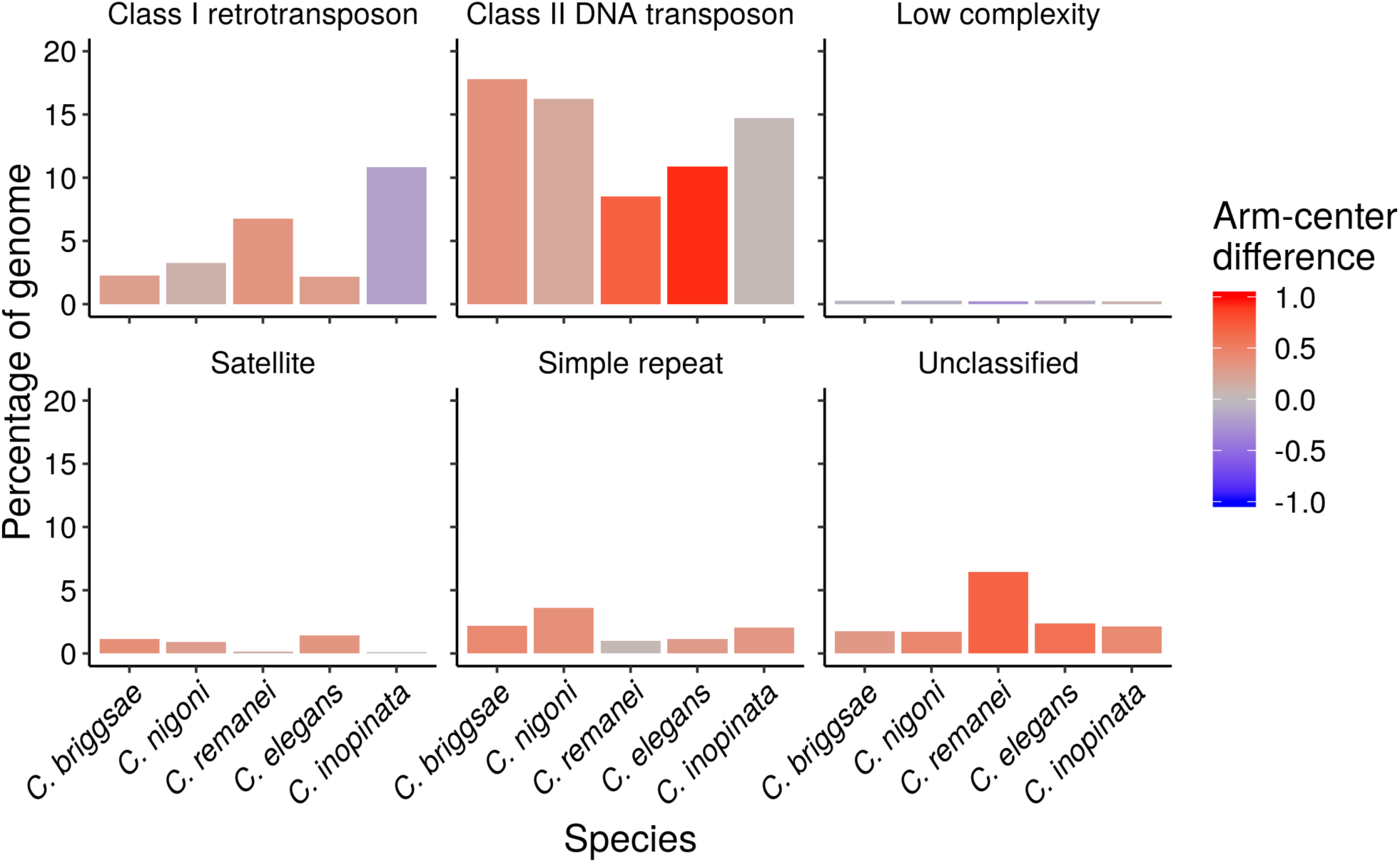
The abundance and genomic landscape of the dominant repeat categories and the two transposable element classes in all species. The y-axis represents the percentage of the genome covered by that repeat category. The color gradient follows the Cohen’s *d* effect size of chromosome position on repeat density as illustrated in Figure 4b. “Low complexity” refers to A-, GA-, or G-rich regions. “Simple repeat” refers to tandem repeats.

**Supplemental Figure 5.**
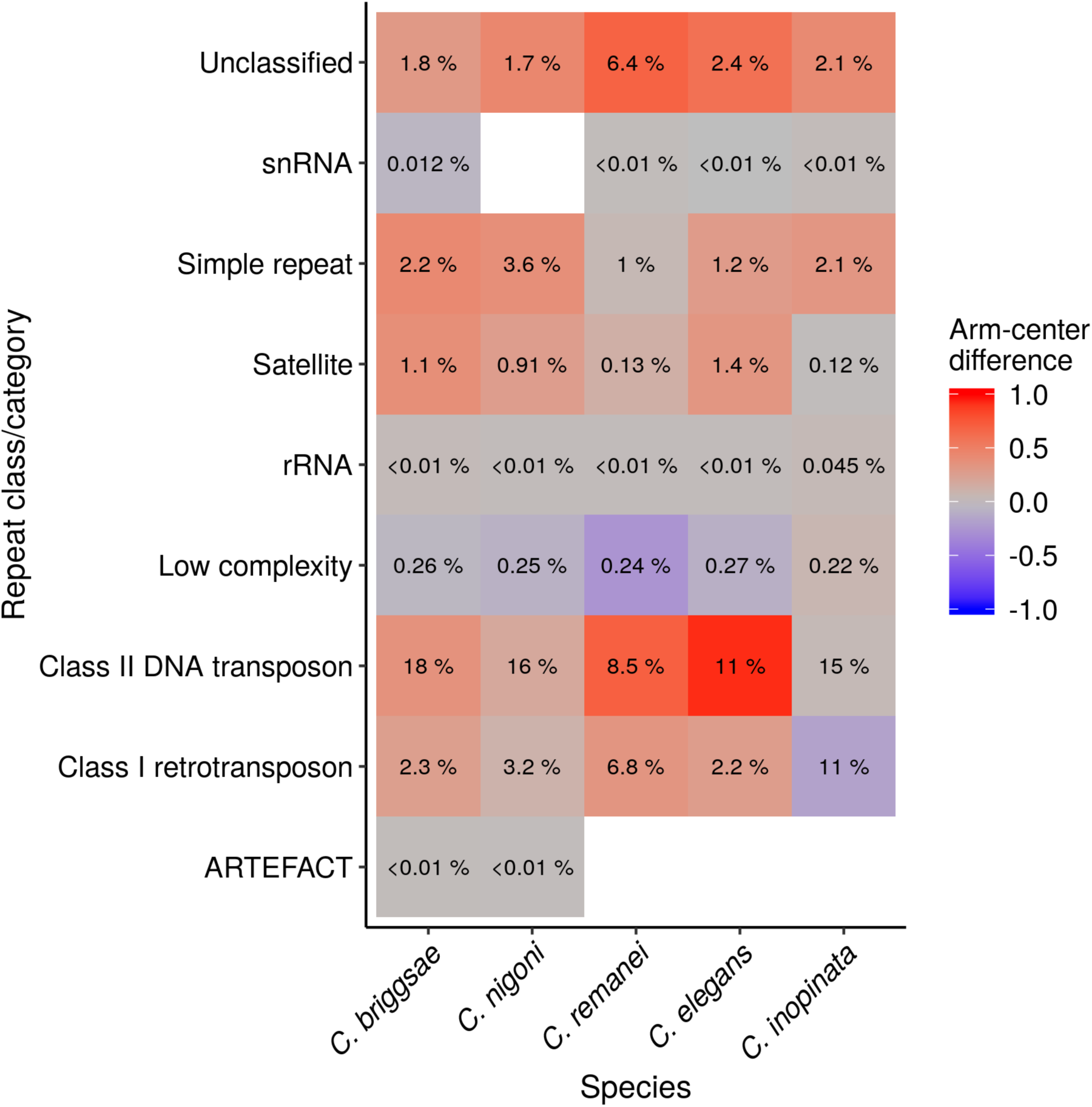
Heat map of the genomic landscape of all repeat categories and the two transposable element classes in all species. The same data as in Supplemental Figure 6 are plotted. The color gradient follows the Cohen’s *d* effect size of chromosome position on repeat density as illustrated in Figure 4b. The numbers in the tiles note the percentage of the genome occupied by the given repeat category. Uncolored tiles represent taxa or categories that were not detected in that species. “Unknown” are repetitive sequences that were not classified by RepeatClassifier. “ARTEFACT” refers to one repetitive element in the *C. briggsae* and *C. nigoni* genomes that aligned to *E. coli* in a BLAST search. “Low complexity” refers to A-, GA-, or G-rich regions. “Simple repeat” refers to tandem repeats.

**Supplemental Figure 6.**
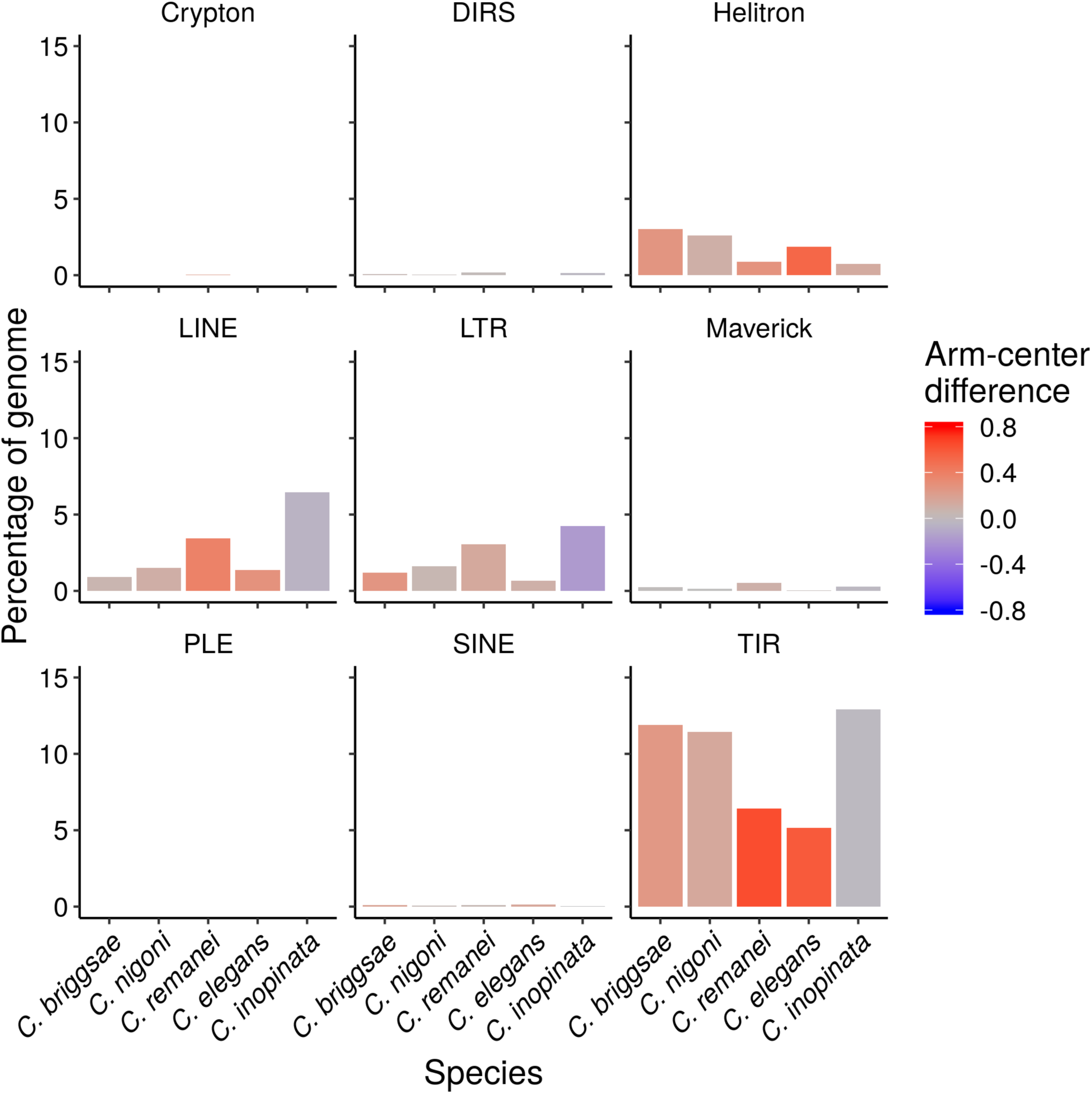
The abundance and genomic landscape of all transposable element orders in all species. The y-axis represents the percentage of the genome covered by that repeat category. The color gradient follows the Cohen’s *d* effect size of chromosome position on repeat density as illustrated in Figure 4b. Some elements in higher taxa were unable to be categorized into lower taxonomic ranks.

**Supplemental Figure 7.**
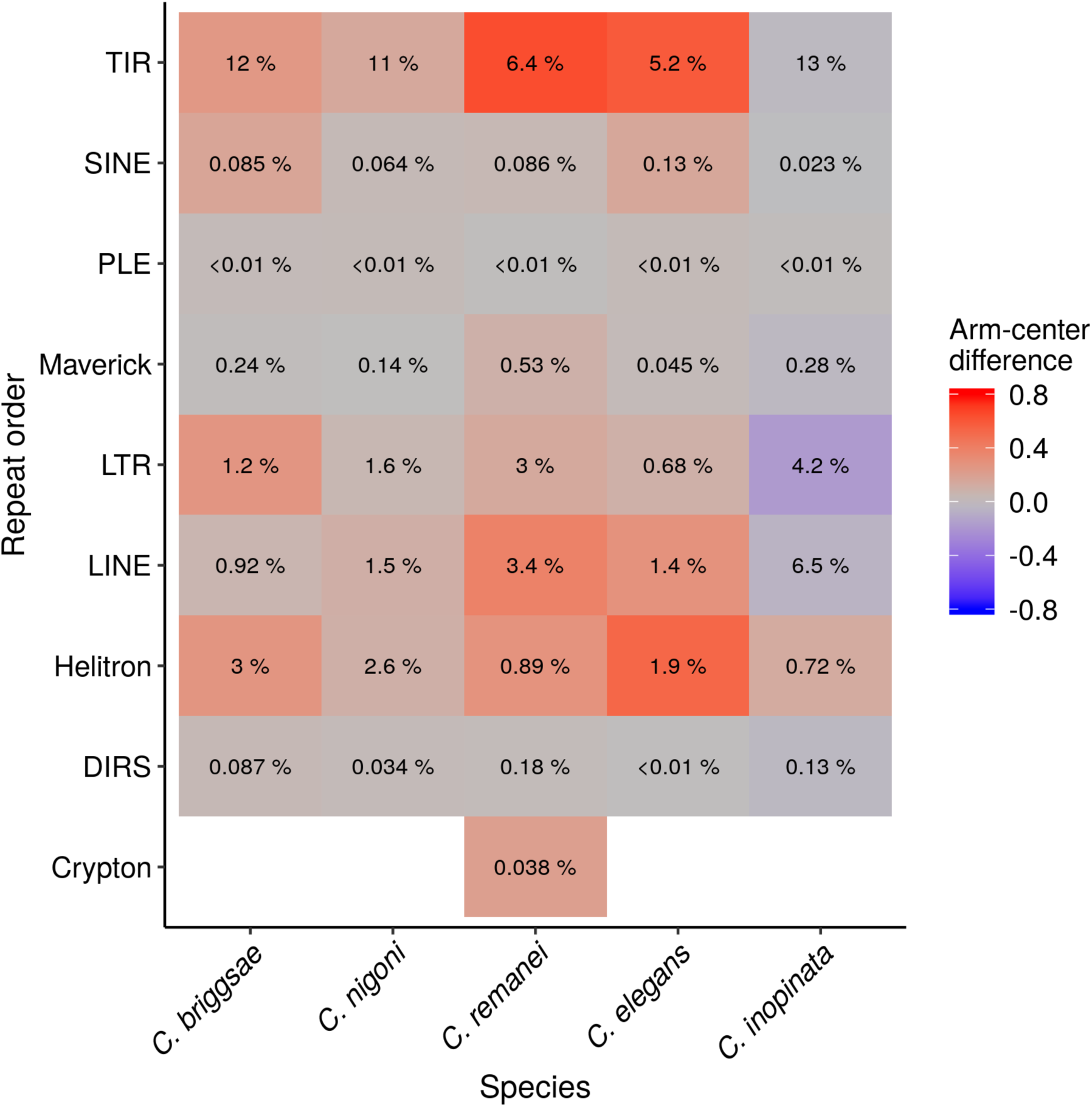
Heat map of the genomic landscape all transposable element orders in all species. The same data as in Supplemental Figure 6 are plotted. The color gradient follows the Cohen’s *d* effect size of chromosome position on repeat density as illustrated in Figure 4b. The numbers in the tiles note the percentage of the genome occupied by the given transposable element order. Uncolored tiles represent taxa that were not detected in that species. Some elements in higher taxa were unable to be categorized into lower taxonomic ranks.

**Supplemental Figure 8.**
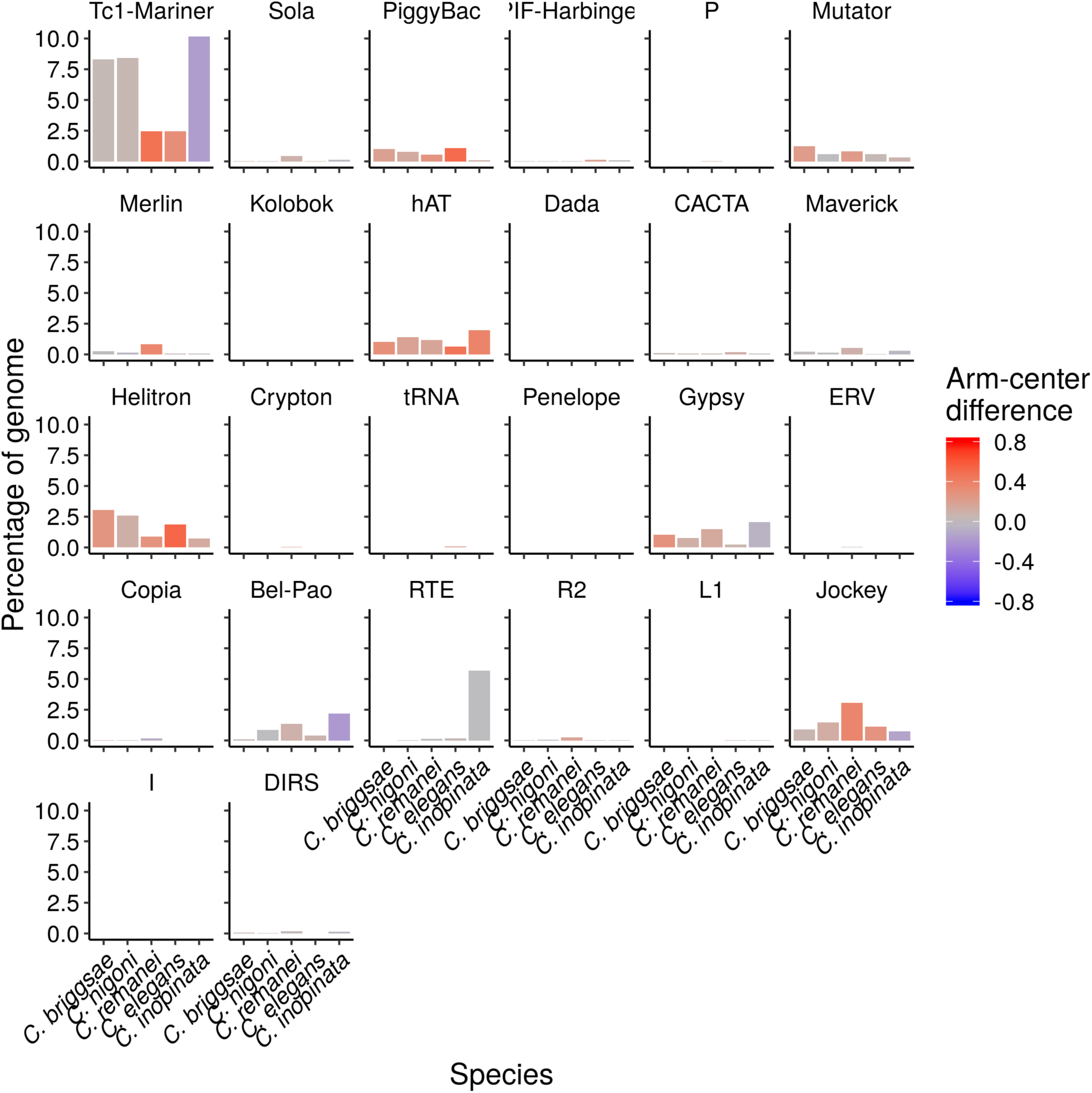
The abundance and genomic landscape of all transposable element superfamilies in all species. The y-axis represents the percentage of the genome covered by that repeat category. The color gradient follows the Cohen’s *d* effect size of chromosome position on repeat density as illustrated in Figure 4b. Some elements in higher taxa were unable to be categorized into lower taxonomic ranks.

**Supplemental Figure 9.**
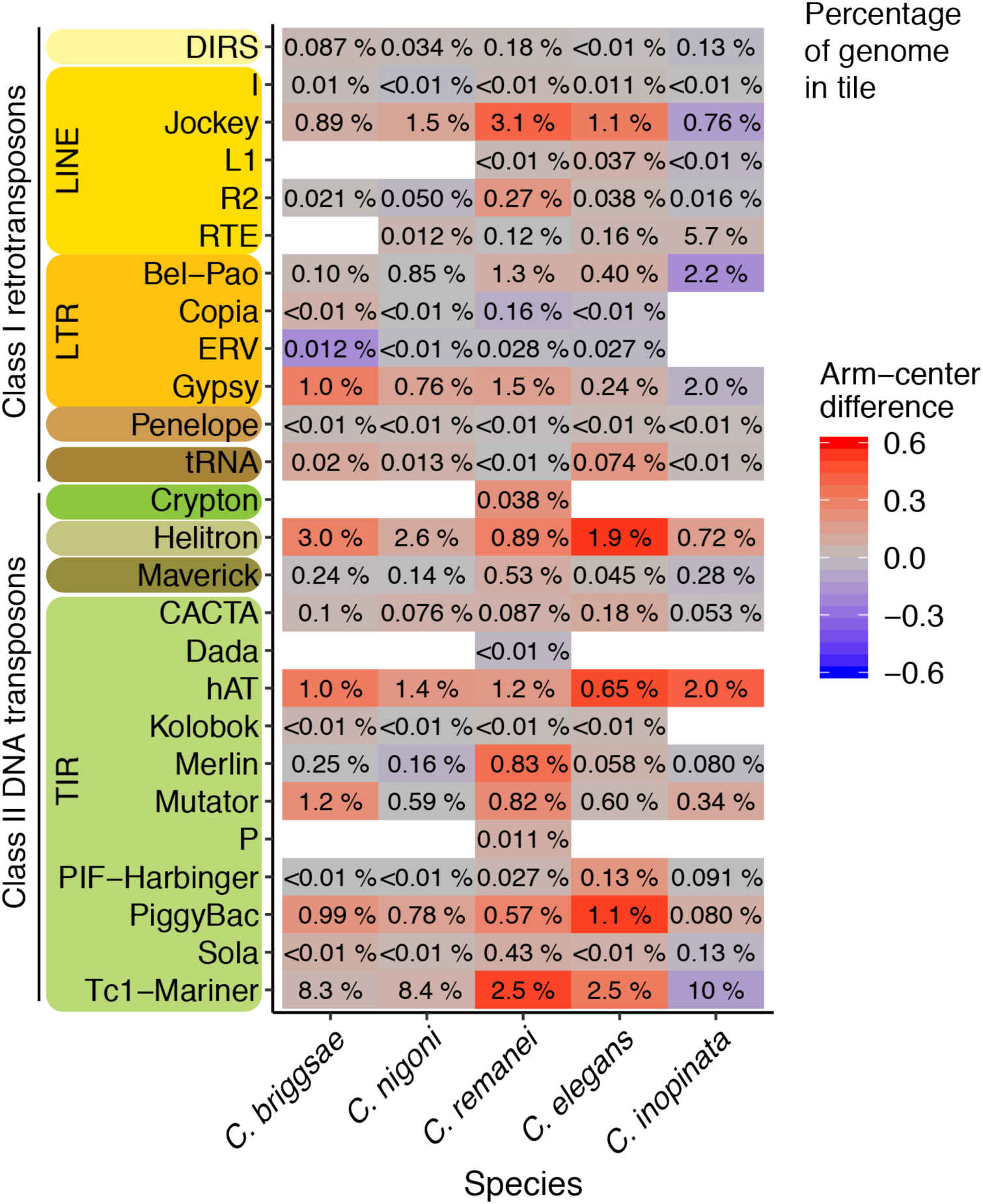
Heat map of the genomic landscape of all transposable element superfamilies in all species. The same data as in Supplemental Figure 8 are plotted. The color gradient follows the Cohen’s *d* effect size of chromosome position on repeat density as illustrated in Figure 4b. The numbers in the tiles note the percentage of the genome occupied by the given repeat superfamily. Superfamilies are clustered by more-encompassing taxonomic groups, and they are colored by repeat orders. Uncolored tiles represent superfamilies that were not detected in that species. Some elements in higher taxa were unable to be categorized into lower taxonomic ranks.

**Supplemental Figure 10.**
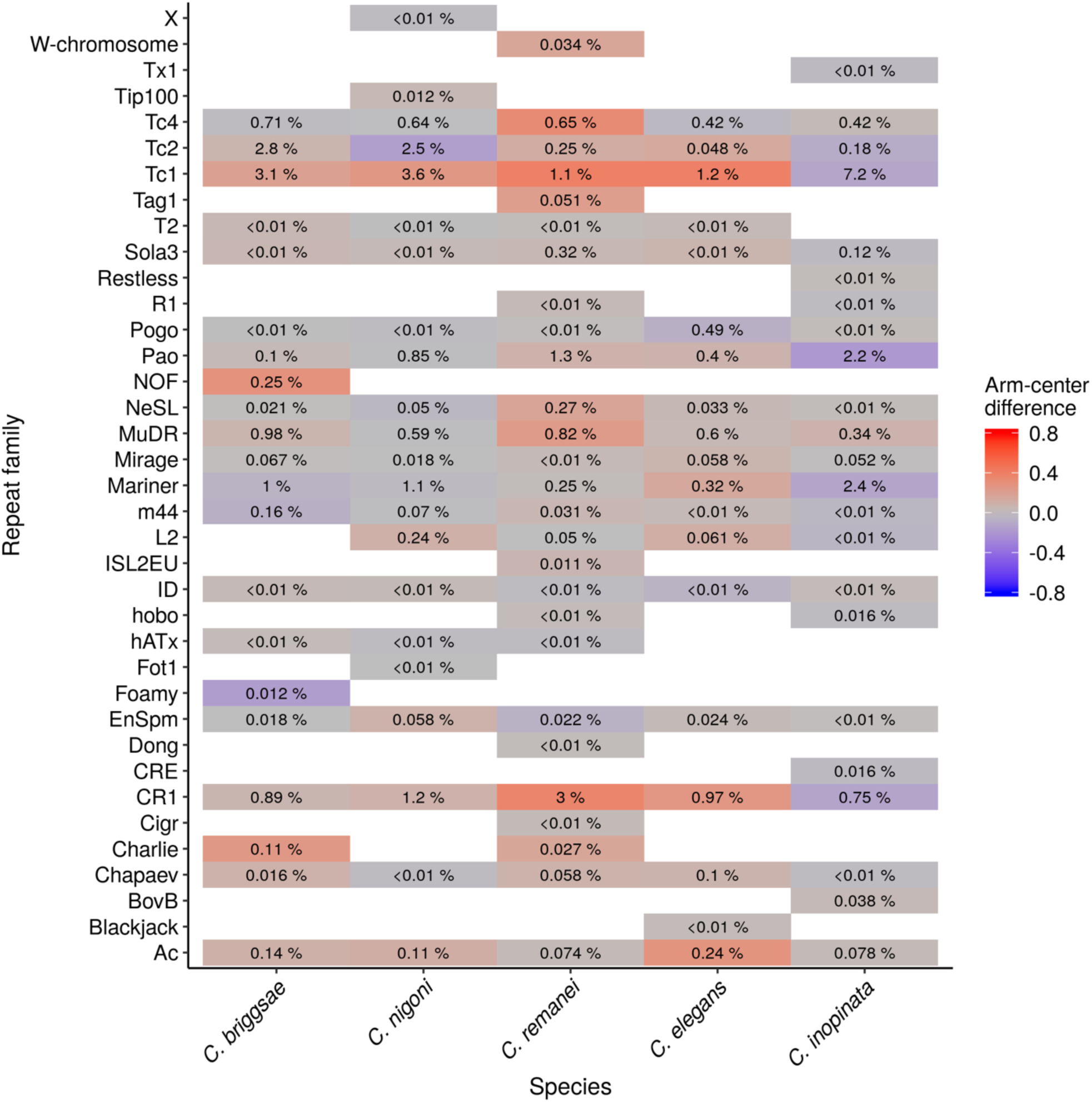
Heat map of the genomic landscape of all transposable element families in all species. The color gradient follows the Cohen’s *d* effect size of chromosome position on repeat density as illustrated in Figure 4b. The numbers in the tiles note the percentage of the genome occupied by the given transposable element family. Uncolored tiles represent families that were not detected in that species. Some elements in higher taxa were unable to be categorized into lower taxonomic ranks.

**Supplemental Figure 11.**
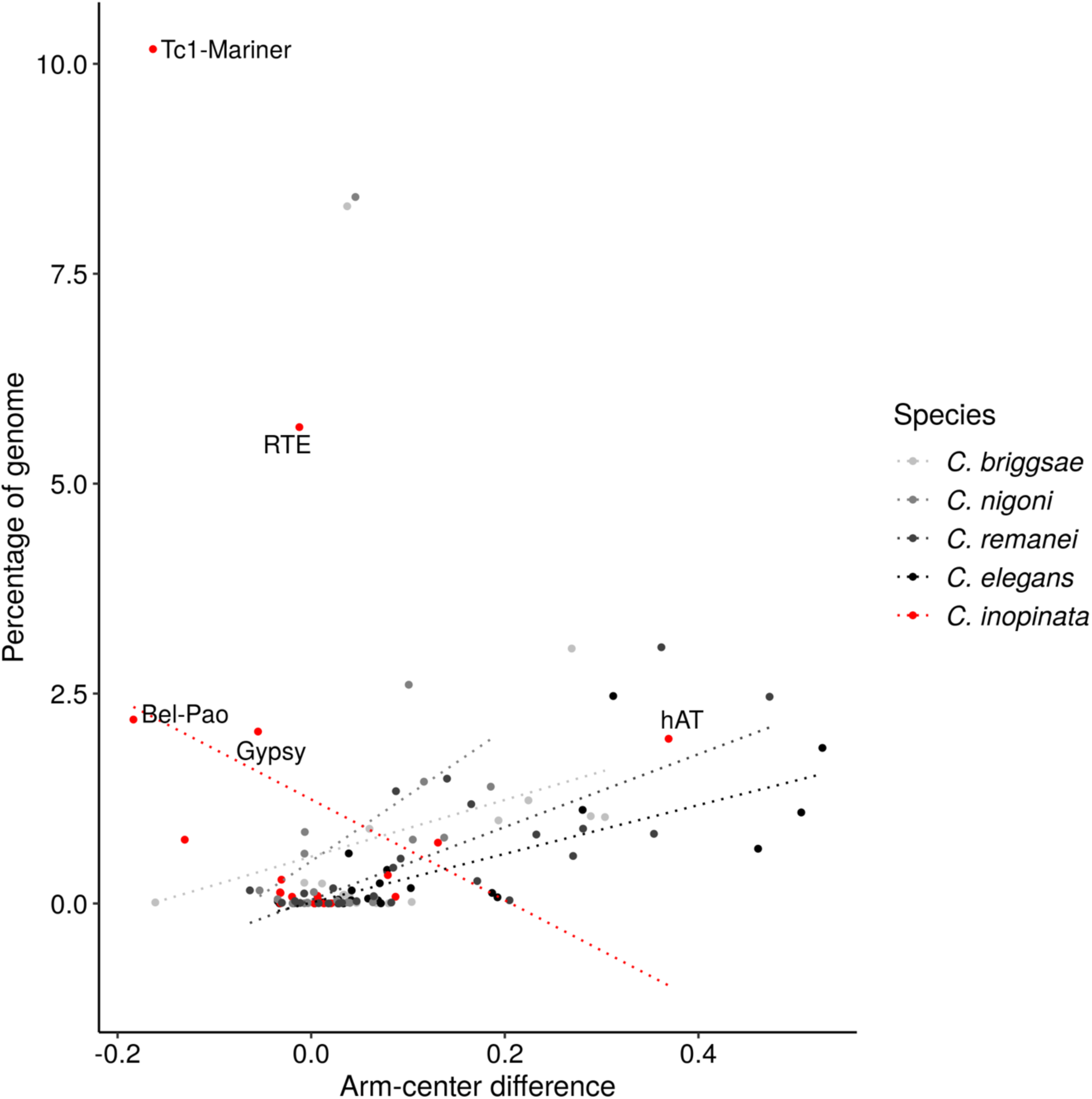
The relationship between repeat chromosomal structure and total genomic repeat content among repeat superfamilies in five *Caenorhabditis* species (not log-transformed). The “arm-center difference” is the Cohen’s *d* effect size of chromosome position on repeat density as described in Figure 4b; this is the same data as Figure 4c but not log-transformed. The five most abundant repeat superfamilies in *C. inopinata* are labeled. Regression statistics can be found in the text and in the supplemental material.

**Supplemental Figure 12.**
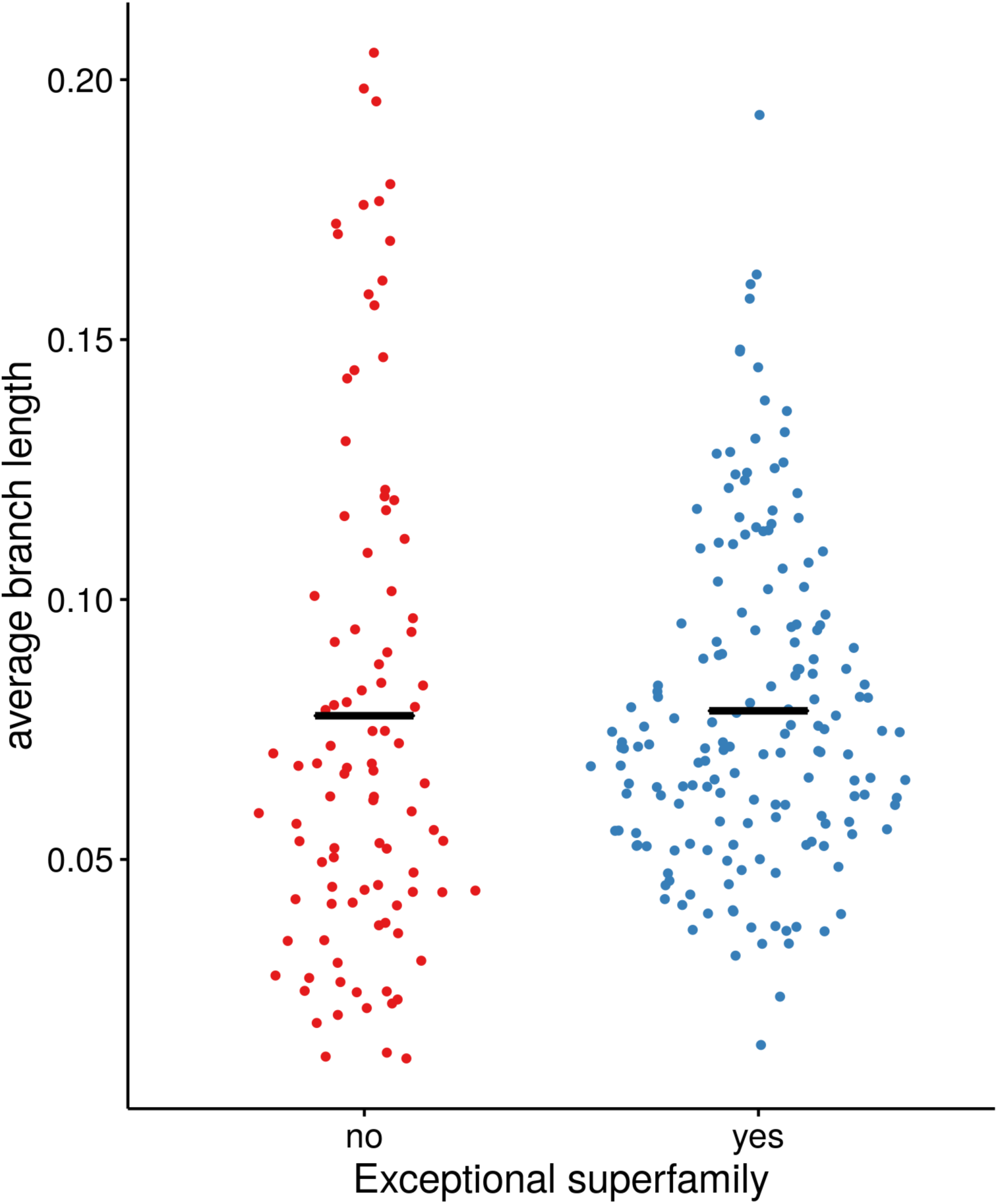
Average branch lengths of repetitive element insertions in *C. inopinata* (untrimmed alignments). Each point represents the mean terminal branch length of a repetitive element cluster phylogenetic tree in *C. inopinata*. Clusters are repetitive elements of the same type as inferred by UCLUST; trees were inferred from alignments of all genomic insertions of that cluster (see methods). Clusters with ≥100 insertions were used in this analysis. The terminal branch lengths of each insertion were used as a measure of divergence. Here, clusters were categorized by whether or not they were in one of the four exceptional repeat superfamilies in *C. inopinata* that are highly abundant and have atypical chromosomal organization (Bel-Pao, Gypsy, RTE, and Tc1-Mariner). Horizontal black line, mean.

**Supplemental Figure 13.**
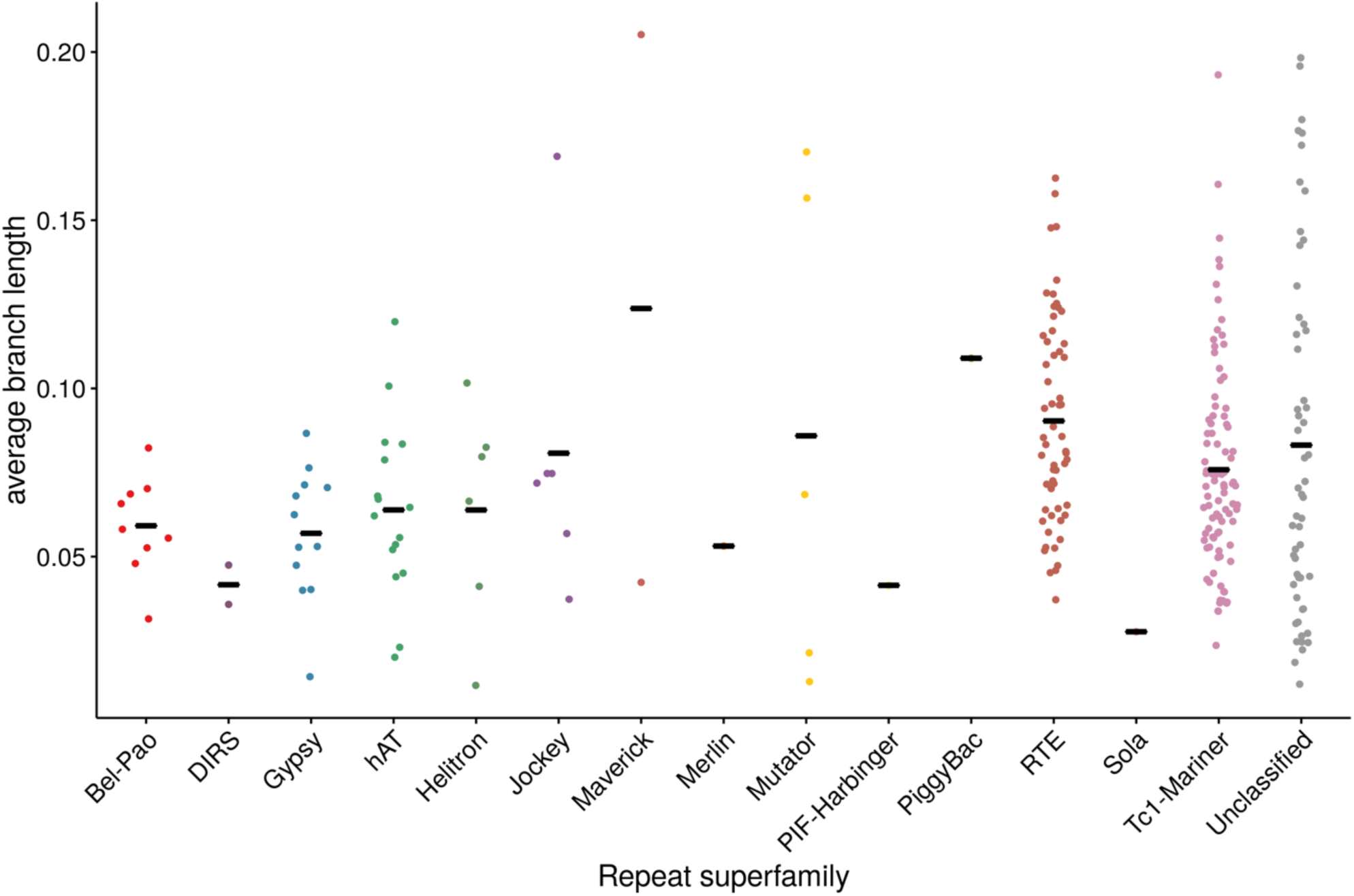
Average branch lengths of repetitive element insertions in *C. inopinata* categorized by repeat superfamily (untrimmed alignments). Each point represents the mean terminal branch length of a repetitive element cluster phylogenetic tree in *C. inopinata*. Clusters are repetitive elements of the same type as inferred by UCLUST; trees were inferred from alignments of all genomic insertions of that cluster (see methods). Clusters with ≥100 insertions were used in this analysis. The terminal branch lengths of each insertion were used as a measure of divergence. Same data as in Supplemental Figure 11. Here, clusters are categorized by repeat superfamily. Black horizontal line, mean.

**Supplemental Figure 14.**
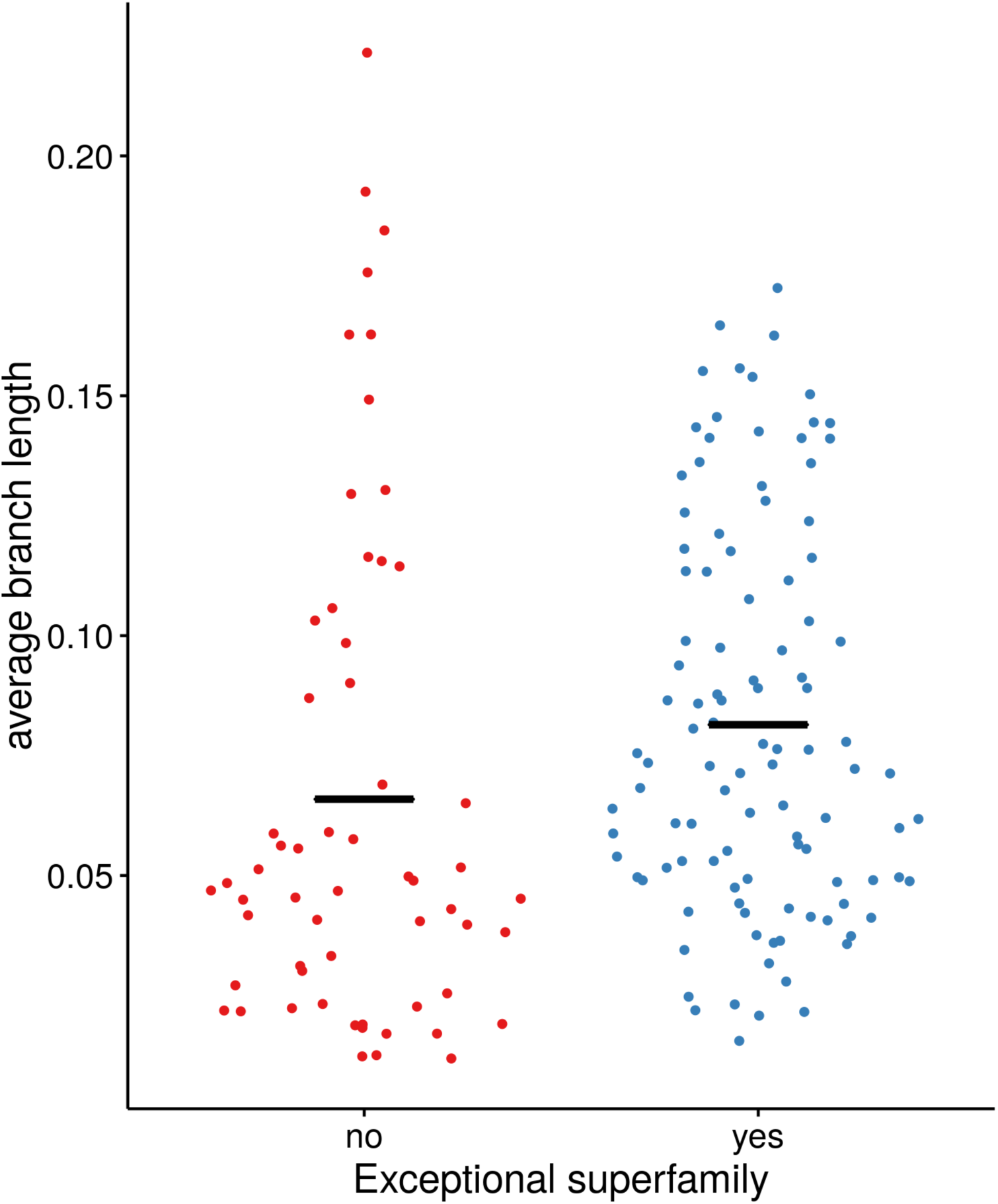
Average branch lengths of repetitive element insertions in *C. inopinata* (trimmed alignments). Each point represents the mean terminal branch length of a repetitive element cluster phylogenetic tree in *C. inopinata*. Clusters are repetitive elements of the same type as inferred by UCLUST; trees were inferred from alignments of all genomic insertions of that cluster (see methods). Clusters with ≥100 insertions were used in this analysis. Trimmed alignments were further filtered to exclude those <50 bp in length; this led to 265 clusters with untrimmed alignments being pared down to 165 clusters with trimmed alignments. The terminal branch lengths of each insertion were used as a measure of divergence. Here, clusters were categorized by whether or not they were in one of the four exceptional repeat superfamilies in *C. inopinata* that are highly abundant and have atypical chromosomal organization (Bel-Pao, Gypsy, RTE, and Tc1-Mariner). Horizontal black line, mean.

**Supplemental Figure 15.**
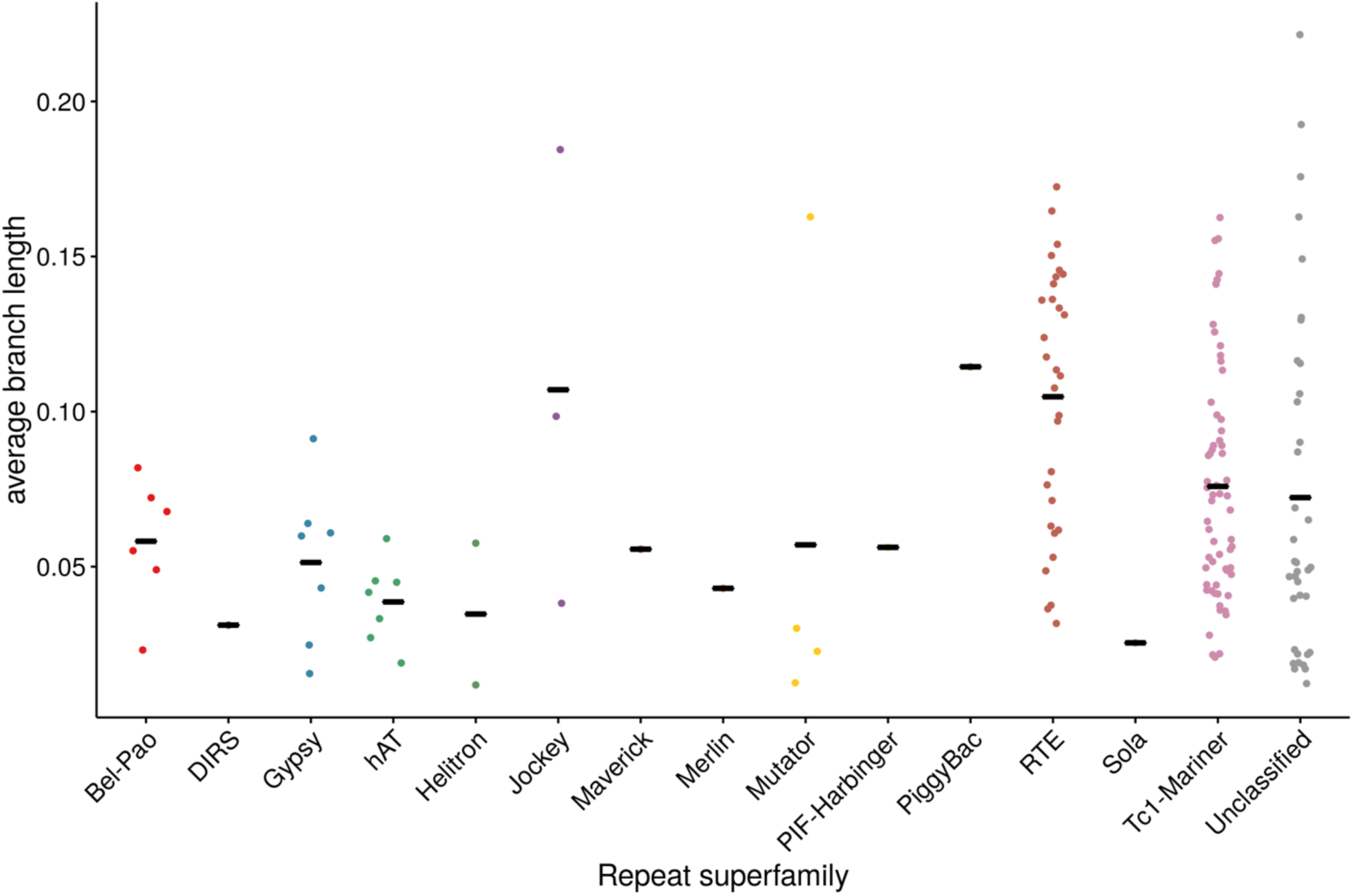
Average branch lengths of repetitive element insertions in *C. inopinata* categorized by repeat superfamily (trimmed alignments). Each point represents the mean terminal branch length of a repetitive element cluster phylogenetic tree in *C. inopinata*. Clusters are repetitive elements of the same type as inferred by UCLUST; trees were inferred from alignments of all genomic insertions of that cluster (see methods). Clusters with ≥100 insertions were used in this analysis. Trimmed alignments were further filtered to exclude those <50 bp in length; this led to 265 clusters with untrimmed alignments being pared down to 165 clusters with trimmed alignments. The terminal branch lengths of each insertion were used as a measure of divergence. Same data as in Supplemental Figure 13. Here, clusters are categorized by repeat superfamily. Black horizontal line, mean.

**Supplemental Figure 16.**
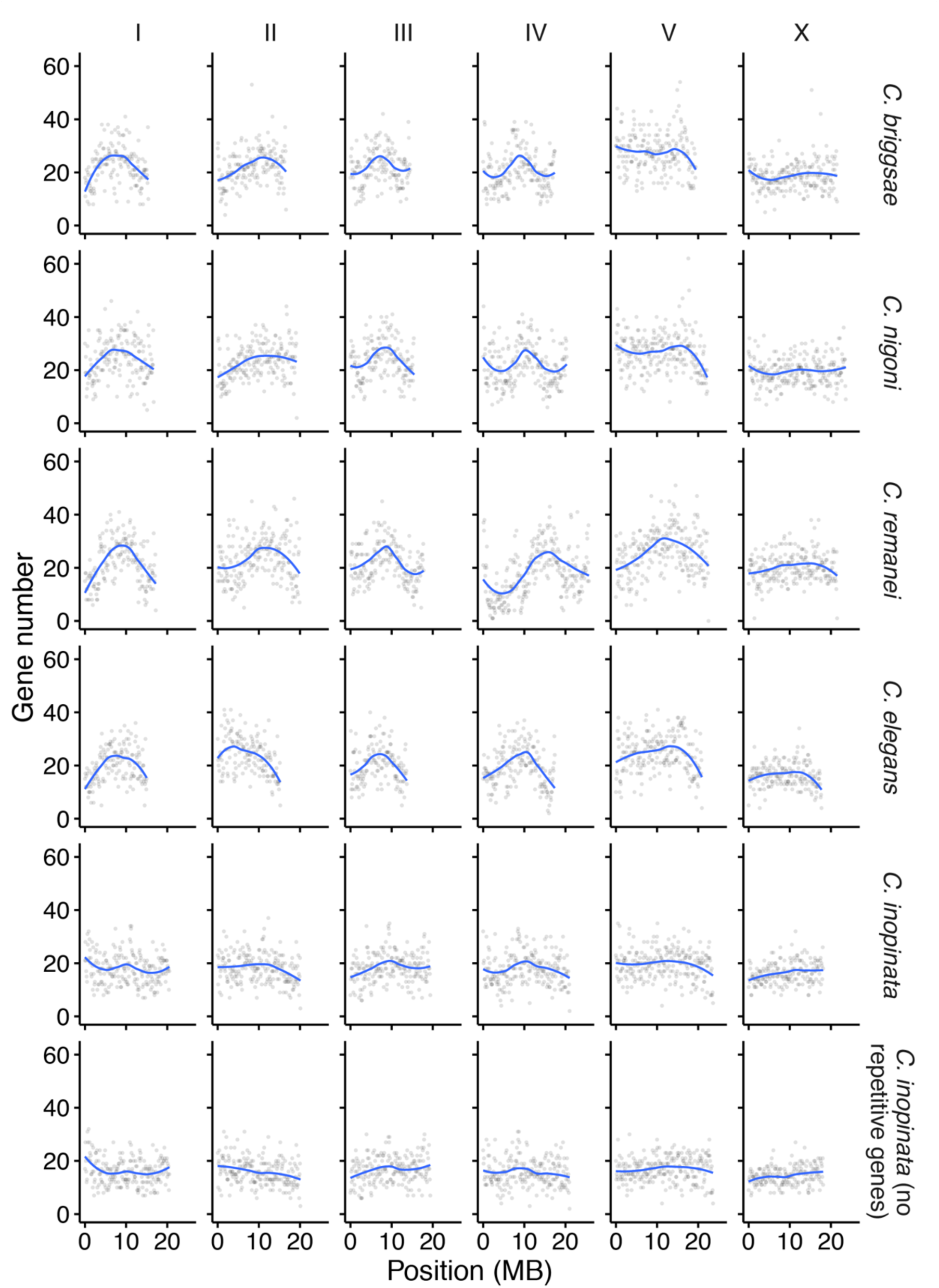
Gene number among 100kb genomic windows in five *Caenorhabditis* species. Here, the gene densities of *C. inopinata* when 2,489 transposon-aligning genes are excluded are also plotted. The blue line was fit by LOESS local regression.

**Supplemental Figure 17.**
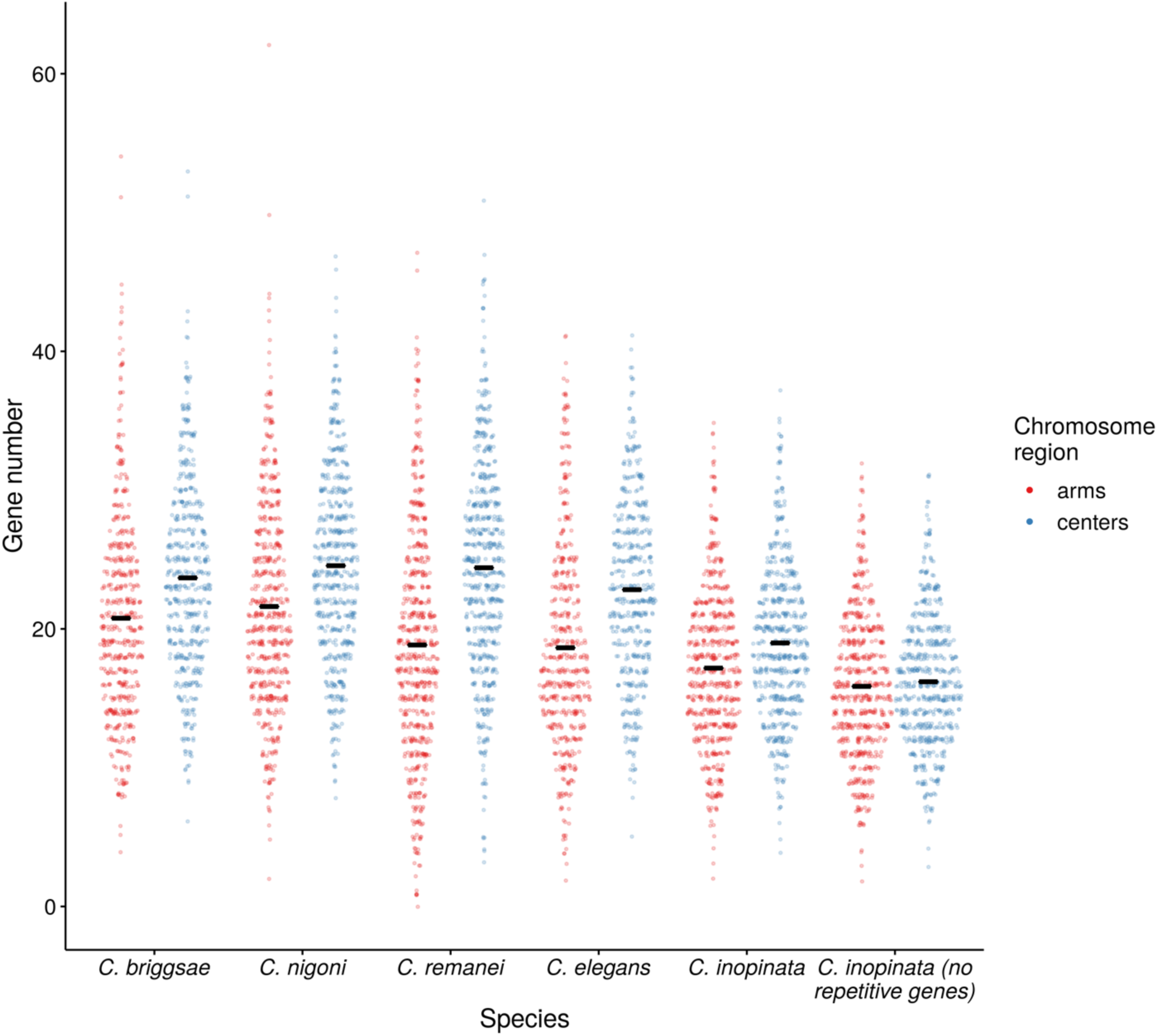
Distribution of gene counts on chromosome arms and centers. Each point represents one 100 kb genomic window. After normalizing chromosome positions by distance to chromosome midpoint, windows were classified as being in chromosome “centers” (middle half of chromosome) or “arms” (outer half of chromosome). Sina plots (strip charts with points taking the contours of a violin plot) reveal the distribution of repeat densities of these windows in chromosome centers and arms for all species. Here, the gene densities of *C. inopinata* when 2,489 transposon-aligning genes are excluded are also plotted (last position on x-axis). Black horizontal lines, means.

**Supplemental Figure 18.**
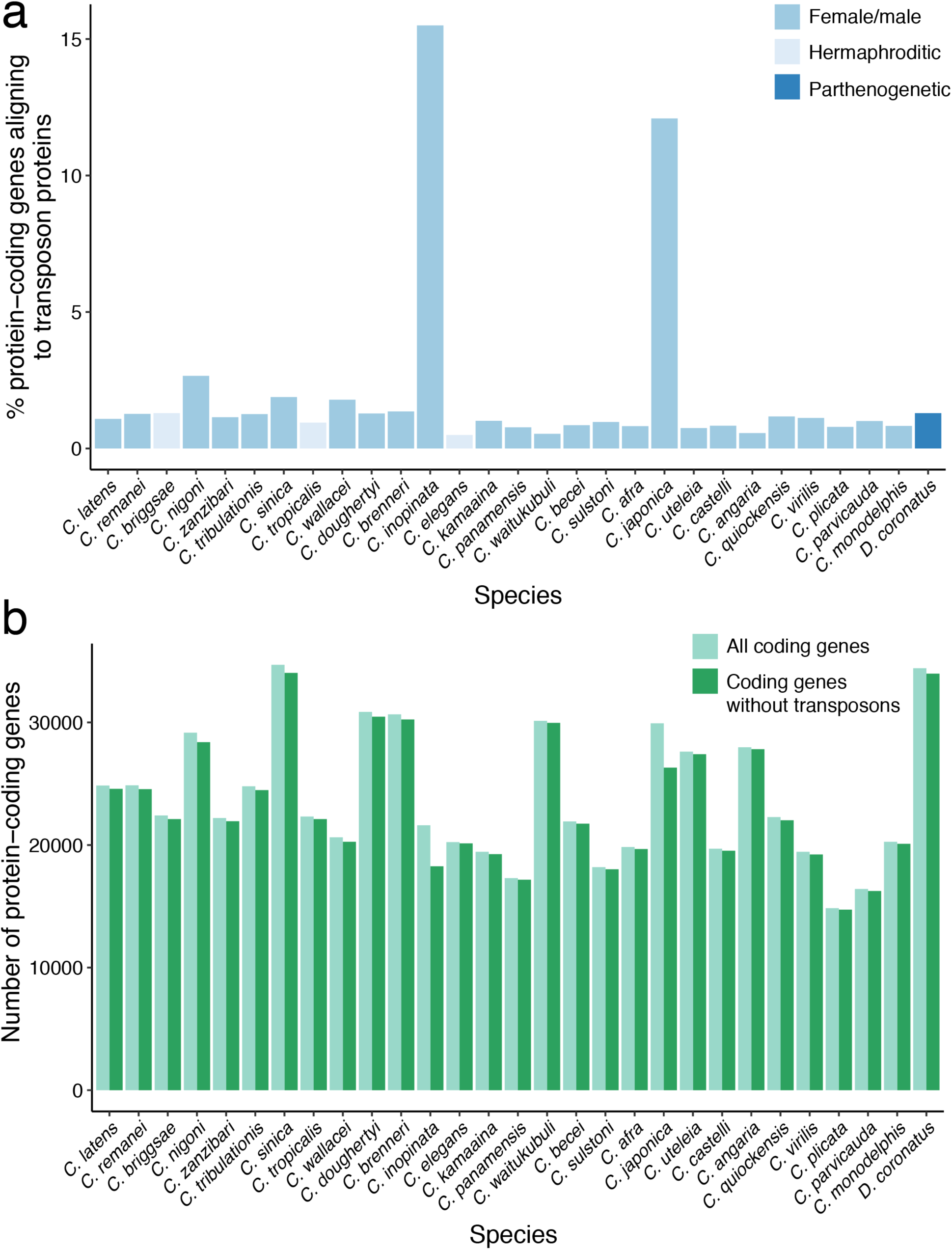
Predicted protein-coding genes in all available *Caenorhabditis* genomes align to transposon-related proteins. The parthenogenetic *Diploscapter coronatus* is included as an outgroup. a. The percentage of protein-coding genes that align to transposons in all available *Caenorhabditis* genomes. Species are roughly ordered by phylogeny and colored by reproductive mode. b. The total number of predicted protein-coding genes before (light green) and after (dark green) transposon-aligning genes are removed.

**Supplemental figure 19.**
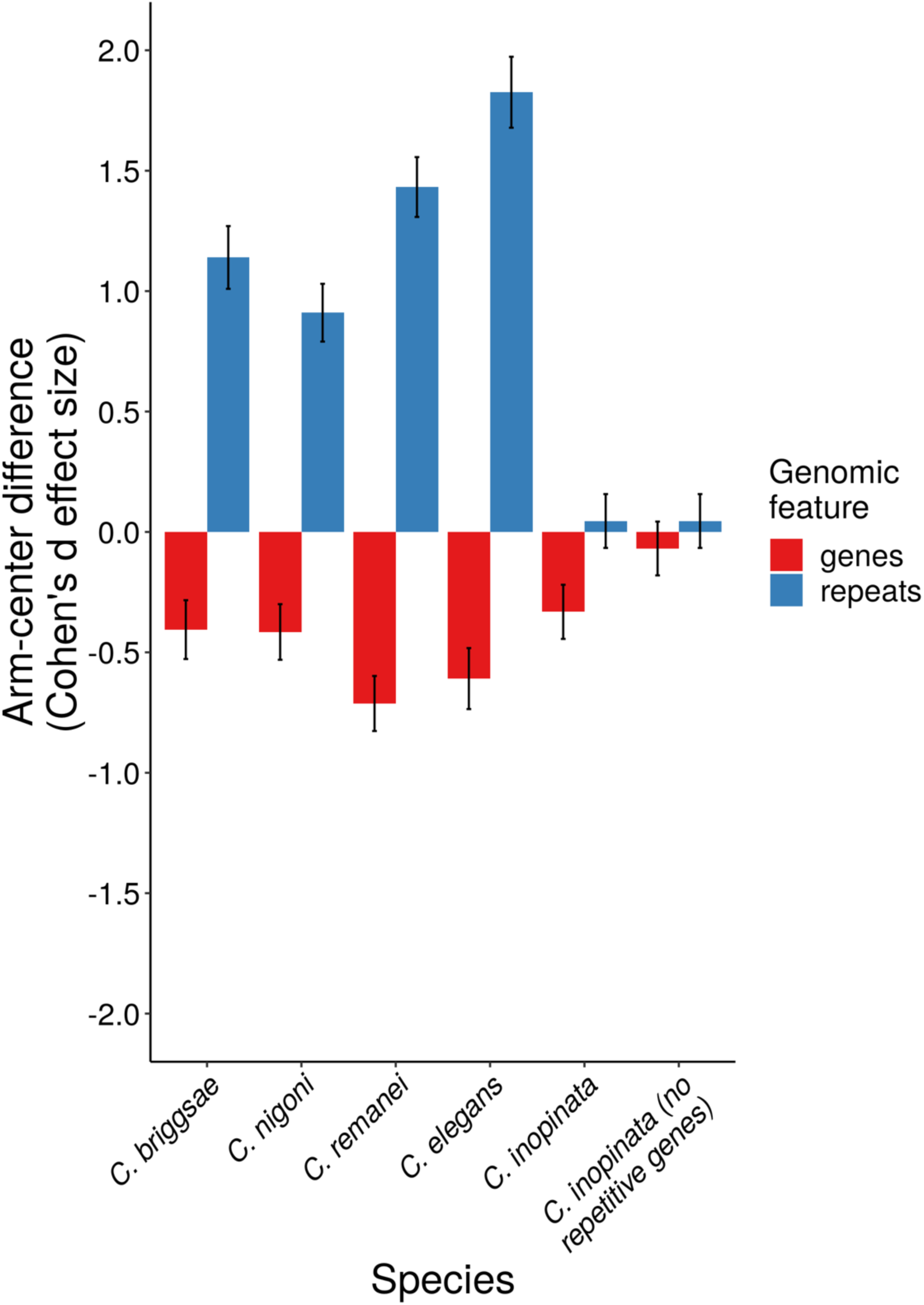
Effect sizes of chromosome position on repeat and gene density. Plotted are the Cohen’s *d* effect sizes of chromosome arms compared to centers on gene and repeat density. Error bars represent 95% confidence intervals. The gene densities of *C. inopinata* when 2,489 transposon-aligning genes are excluded are also plotted (last position on x-axis). These effect sizes are based on repeat percentages and gene counts in 100 kb genomic windows (same data as Figure 7).

**Supplemental figure 20.**
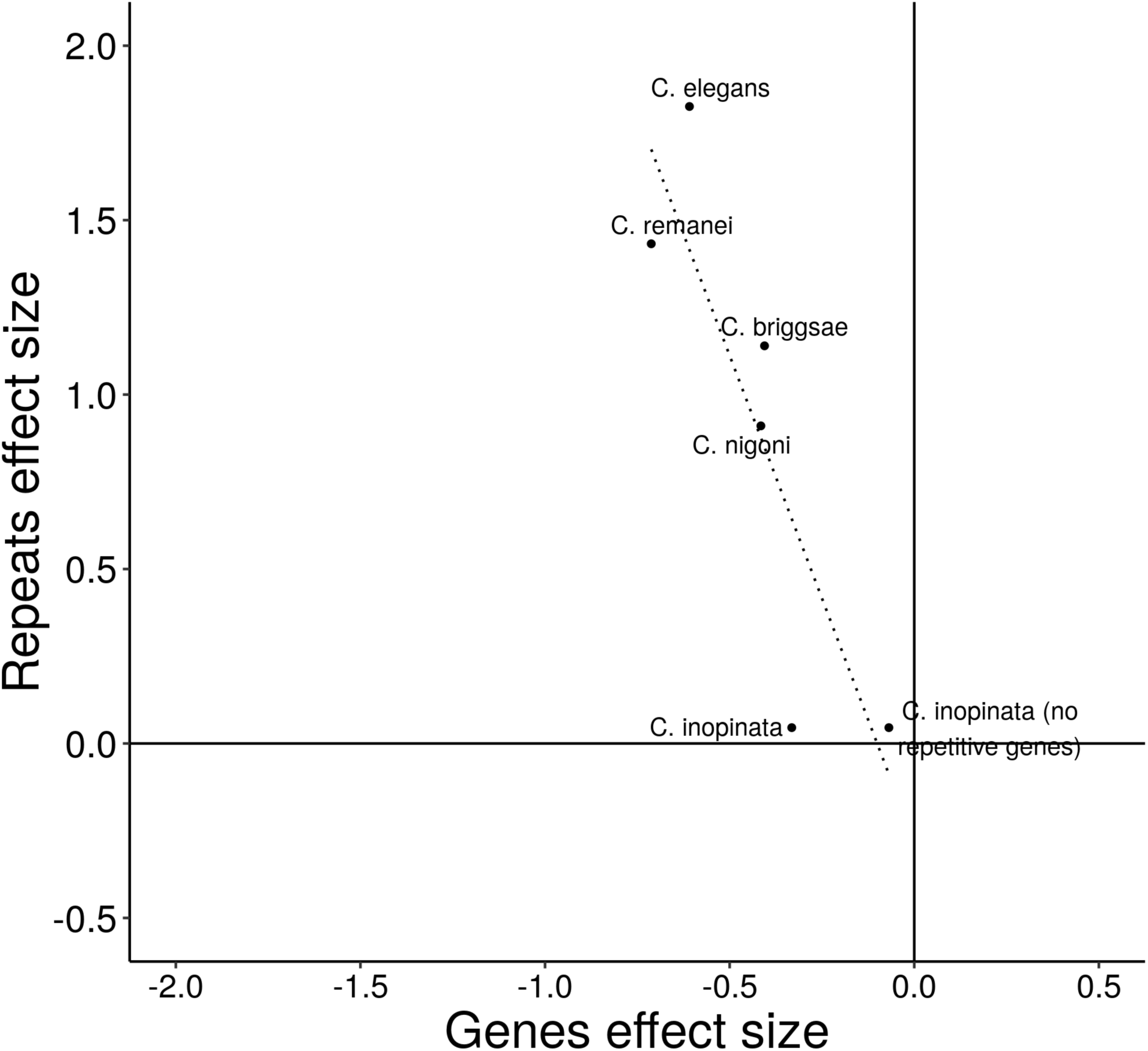
Effect sizes of chromosome position on repeat density covaries with effect sizes of chromosome position on gene density. Plotted are the Cohen’s *d* effect sizes of chromosome arms compared to centers on gene and repeat density. The gene densities of *C. inopinata* when 2,489 transposon-aligning genes are excluded are also plotted. These effect sizes are based on repeat percentages and gene counts in 100 kb genomic windows (same data as Figure 7 and Supplemental figure 18). The linear fit is significant (r^2^=0.67; *F*=11; β_1_= −2.8; *p*=0.029) but dubious as *C. inopinata* is double-counted and the sample is small. When *C. inopinata* (including transposon-aligning genes) is excluded, the fit remains significant (r^2^=0.81; *F*=18; β_1_= −2.5; *p*=0.024).

**Supplemental figure 21.**
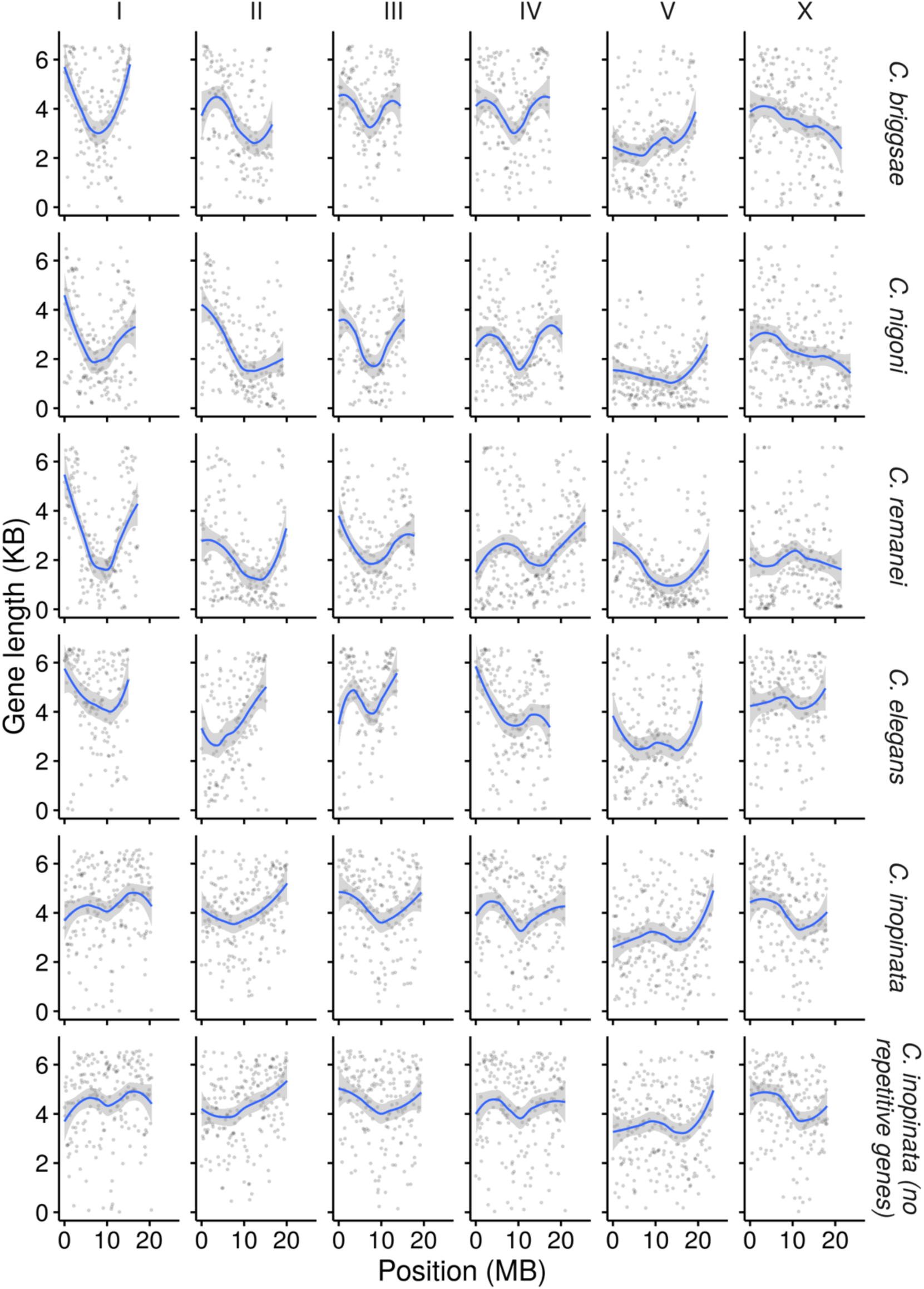
Gene length among 100kb genomic windows in five *Caenorhabditis* species. Plotted is the average gene length in that window. Here, the genes of *C. inopinata* when 2,489 transposon-aligning genes are excluded are also plotted. The blue line was fit by LOESS local regression; the shaded ribbon is the 95% confidence interval.

**Supplemental figure 22.**
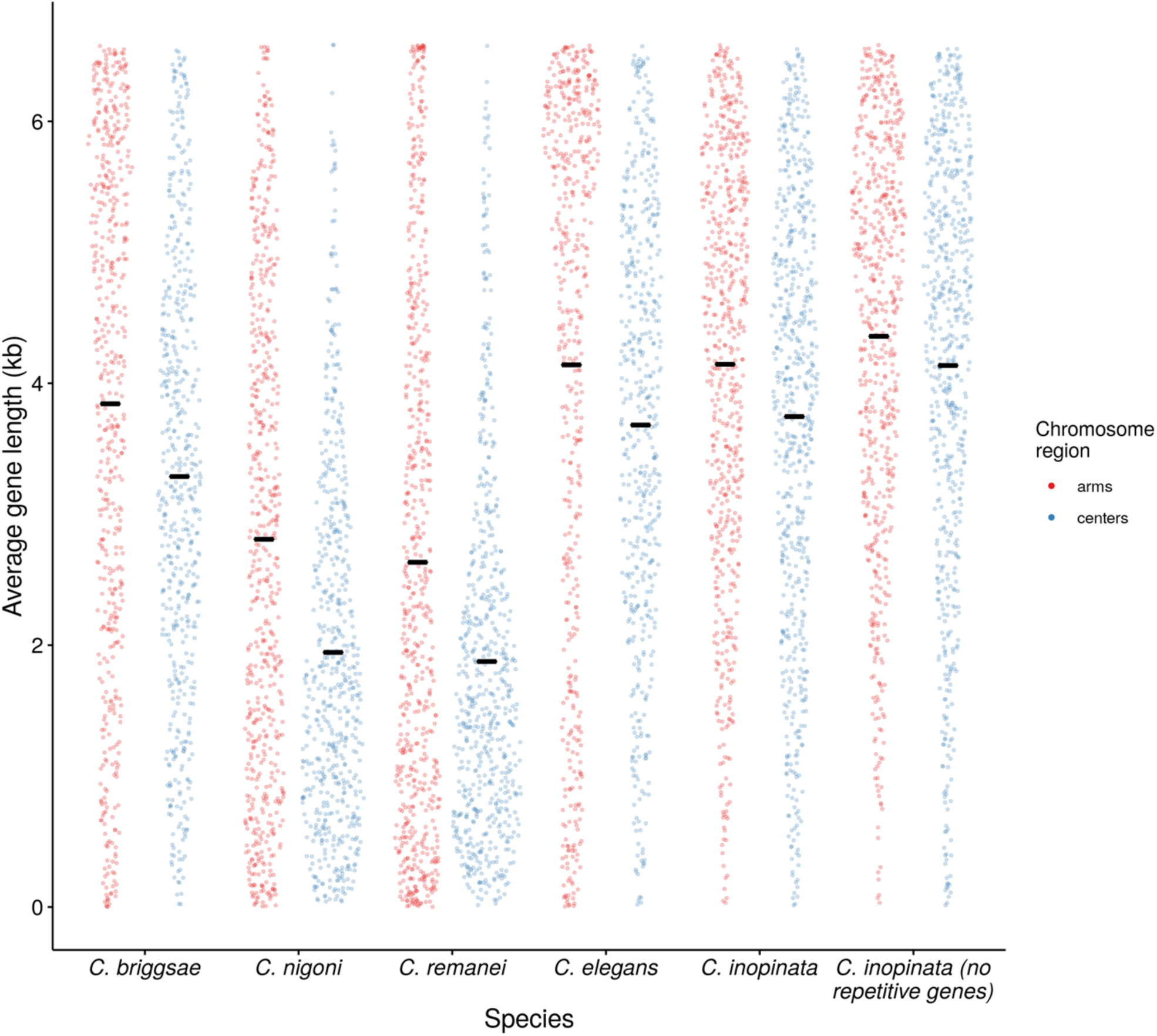
Distribution of gene lengths on chromosome arms and centers. Each point represents one 100 kb genomic window and the average gene length in that window. After normalizing chromosome positions by distance to chromosome midpoint, windows were classified as being in chromosome “centers” (middle half of chromosome) or “arms” (outer half of chromosome). Sina plots (strip charts with points taking the contours of a violin plot) reveal the distribution of repeat densities of these windows in chromosome centers and arms for all species. Here, the gene densities of *C. inopinata* when 2,489 transposon-aligning genes are excluded are also plotted (last position on x-axis). Black horizontal lines, means.

Supplemental data (statistics_summary.xls; repeat_classes_genomic_landscapes.pdf; repeat_orders_genomic_landscapes.pdf; repeat_superfamilies_genomic_landscapes.pdf; repeat_families_genomic_landscapes.pdf) are available at *Journal*. Data, code, and additional supplemental figures associated with this study have been deposited in Github at https://github.com/gcwoodruff/transposable_elements_2019.

## Data deposition and accessibility

Genome sequences and annotations were retrieved from WormBase ParaSite (parasite.wormbase.org) and the *Caenorhabditis* Genomes Project (Caenorhabditis.org). Data, code, and additional supplemental figures associated with this study have been deposited in Github at https://github.com/gcwoodruff/transposable_elements_2019.

## Author contributions

GCW re-masked genome assemblies, analyzed data, made figures, and wrote the first draft of the paper. AAT performed evolutionary simulations and informed the masking approach. GCW and AAT revised the paper.

## Acknowledgements

We thank Patrick Phillips for sharing genomic data. Patrick Phillips, Bill Cresko, Peter Ralph, Andy Kern, and their laboratory members provided helpful comments throughout the development of this work. Jeffrey Adrion provided valuable feedback on earlier versions of this manuscript. This work was supported by funding from the National Institutes of health to GCW (5F32GM115209-03) and to Patrick Phillips (R01 GM-102511; R01 AG049396).

